# Pooled progeny sequencing effectively estimates polyandry in species with large families

**DOI:** 10.64898/2026.10.02.756305

**Authors:** Todd A. Farmer, Ashley Bentz, Stuart Macdonald, Robert L. Unckless, John K. Kelly

## Abstract

Polyandry, multiple mating by females, both benefits females by increases genetic variation within families and provides an opportunity for sexual selection through sperm competition and cryptic female choice. Estimating polyandry in species with large clutches is challenging because genotyping all progeny individually can be impractical. We present a likelihood-based framework for estimating the number of sires and their relative contributions from the maternal genotype and pooled sequencing of entire progeny pools. The method evaluates the probability of observed pooled offspring read counts under alternative paternal genotype mixtures, accounting for Mendelian segregation and sampling variation from finite sequencing depth. We implement this framework for both biallelic SNPs and short-read mini-haplotype markers constructed from closely linked SNPs, which provide multi-allelic information comparable to traditional microsatellite markers. Simulations show that this approach accurately recovers sire number and sire contributions across a range of conditions. We then applied the method to 275 wild-caught *Drosophila melanogaster* females and their offspring from a Kansas population using 3,053 biallelic SNPs and 239 four-allele mini-haplotype markers. Estimates from the two marker sets were highly concordant, and combined analyses indicate that 57% of families were sired by a single male. These results demonstrate that pooled progeny sequencing provides an efficient genomic approach for estimating polyandry in species with large families.

## Introduction

Polyandry, defined as female reproduction involving multiple males within a reproductive period, is widespread across animals and plants and has important evolutionary consequences (Jennions & Petrie, 2000; Simmons, 2005; Collet et al., 2012; Pannell & Labouche, 2013). By increasing genetic diversity within families, polyandry may broaden the distribution of offspring fitness and buffer maternal reproductive success against environmental heterogeneity, parasites, pathogens, or genetically incompatible mates (Jennions & Petrie, 2000; Slatyer et al., 2011; Holman, 2015). With polyandry, competition between males to fertilize the limited number of eggs may drive selection for complex male reproductive anatomy and accessory gland proteins that alter female behavior or physiology, to increase a male’s share of paternity (Clark & Begun, 1998; Fiumera et al., 2007). In turn, females may counter-adapt to maintain control over their own reproductive output in the face of manipulative male traits. This conflict between the sexes can shift the arena of sexual selection from pre-copulatory traits, such as courtship displays, to post-copulatory physiological adaptations, with potential consequences for speciation and phenotypic divergence (Birkhead & Pizzari, 2002; Kvarnemo & Simmons, 2013).

Quantifying polyandry requires estimating both the number of males that sire offspring within a female’s clutch and the relative contribution of each male (Jones et al., 2009; Turnell & Shaw, 2015). The distribution of paternal contributions is important for understanding the strength of sperm competition and the opportunity for post-copulatory sexual and gametic selection (Kvarnemo & Simmons, 2013; Wedell, 2013; Turnell & Shaw, 2015). The outcome, the distribution of paternal contributions among zygotes, determines how genetic diversity is partitioned within families and the relatedness structure of maternal sibships. This, in turn, determines the efficacy of evolutionary processes such as kin selection (Boomsma & Ratnieks, 1996; Hughes et al., 2008).

A common approach to estimating polyandry is to genotype mothers and individual offspring and then infer the number of paternal genotypes required to explain the observed progeny genotypes (Jones, 2005; Eriksson et al., 2009; Colicchio et al. 2020). This strategy has been especially successful when highly polymorphic multi-allelic markers, such as allozymes or microsatellites, are utilized (Ochando et al., 1996; Harshman & Clark, 1998; Imhof et al., 1998; Good et al., 2006). At a given locus, a clutch containing more non-maternal alleles than can be explained by a single diploid sire provides strong evidence for multiple paternity. However, genotyping individual progeny in systems where a single female may produce hundreds to thousands of offspring may not be feasible (Eriksson et al., 2009). Genotyping only a random subsample of offspring may distort estimates of paternal contribution, failing to detect minor sires when paternal contributions are highly skewed (Quennessen et al., 2025).

Pooled sequencing of entire clutches offers a potential solution to this problem. Rather than sequencing offspring individually, all progeny from a maternal family can be combined into a single DNA pool and sequenced together (Schlötterer et al., 2014; Guirao-Rico & Gonzalez, 2021). This approach retains information from the entire clutch and avoids the potential bias associated with progeny subsampling (Quennessen et al., 2025). However, pooled sequencing introduces an inferential problem: The data are read counts for alleles in the pooled offspring sample which must be interpreted in relation to the known genotype of the mother. Estimating the relative contribution to progeny from each male from such data requires a model that predicts the likelihood of pooled progeny read counts under alternative paternal contribution scenarios.

Here, we develop a likelihood model for estimating polyandry from pooled sequencing of maternal families combined with individual genotyping of mothers (Fig 1). We use the language of polyandry, but the model is equally applicable to polygyny in species where the known parent is male (e.g. pipefish; Flanagan et al., 2017). For each family, the maternal genotype is known while the paternal genotype or genotypes are unknown. The model predicts the likelihood of the observed maternal and clutch genomic data as a function of the number of sires and their relative contributions. Because the mother contributes half of the genome of every offspring, her genotype strongly constrains the composition of the offspring pool. The remaining signal reflects the unknown paternal contribution. Candidate models with one sire, two sires, three sires, and so on generate different expected allele frequency patterns in the pooled progeny. The likelihood framework compares these expectations to the observed read counts while accounting for two sequential sources of sampling variation, genetic segregation among offspring and read sampling at a finite sequencing depth.

**Figure 1.**
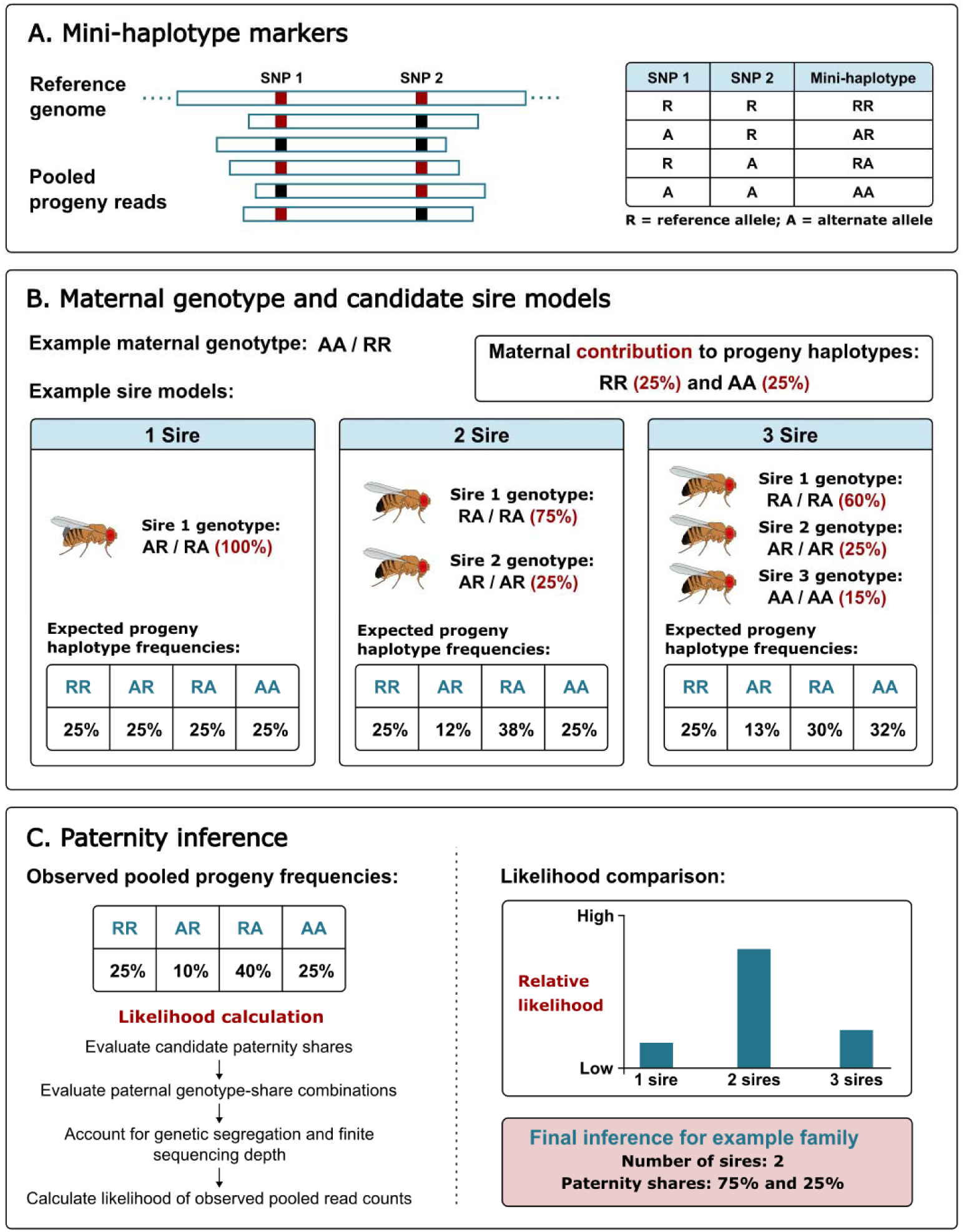
Pooled-sequencing framework for estimating multiple paternity using mini-haplotype markers. A) Four-allele mini-haplotype markers are constructed from pairs of closely linked SNPs that can be observed on the same sequencing read. B) Candidate models differing in sire number, paternal genotypes, and relative paternal contributions are evaluated over possible ordered combinations. Example genotype and paternity-share configurations are shown for one-, two-, and three-sire models. C) For each candidate model, the likelihood of the observed pooled progeny read counts is calculated for all possible paternal genotypes. Relative support among candidate models is then used to infer the number of sires and their estimated proportional contributions for each family.

On a per-marker basis, biallelic Single Nucleotide Polymorphisms (SNPs) are less informative than multi-allelic markers for paternity inference (Hauser et al., 2011; Flanagan & Jones, 2019). We therefore also construct short-read mini-haplotypes from closely linked SNPs (Fig 1A). Closely linked SNPs will frequently be covered by the same sequencing read or read-pair. Coding the two alleles at each SNP as R and A, a read traversing two SNPs can be assigned to one of four possible haplotype states: RR, RA, AR, or AA. These four “mini-haplotypes” can then be treated as allelic states of a single marker (Monnahan et al., 2021; Baetscher et al., 2017). With close linkage, recombination should “break” these alleles very rarely in the single round of meiosis from parent to offspring. This procedure can be extended to encompass more than two closely linked SNPs; there are up to 8 mini-haplotypes with 3 SNPs and 16 with 4 SNPs. Of course, closely linked SNPs often exhibit linkage disequilibria and therefore not all possible haplotypes may be evident in the data (or even present in the population). The advantage of genomic data is scale: With many sets of closely linked loci to choose from, researchers can select a subset that is most informative for estimation.

We implement this theory in Python programs that estimate sire number and paternal contributions from bi- and/or multi-allelic markers. We first evaluate the performance of the method using simulations in which the true number of sires and their respective contributions are known. These simulations demonstrate the accuracy of inference under different sire-contribution scenarios, marker types, and experimental designs (number of families, size of families, sequencing depth). We then apply the method to a field study of wild caught, field-mated *Drosophila melanogaster* females from a Kansas population. Previous studies of wild *Drosophila* using allozymes and microsatellite markers have demonstrated multiple paternity in natural populations (Ochando et al., 1996; Imhof et al., 1998; Good et al., 2006). We extend polyandry estimation to pooled genomic sequencing of entire clutches. We first describe the theory to estimate and characterize polyandry, then the simulation algorithms used to test this theory, followed by the experimental application to a wild population of *Drosophila melanogaster*. The application involves both SNP and mini-haplotype markers and the final estimates synthesize both sets of markers.

## Materials and Methods

### Theory

We calculate the probability of the observed maternal genotype and pooled clutch genomic data as a function of sire number and relative paternal contribution. We assume that the maternal genotype is correctly determined from the data. For an autosomal marker with alleles *A*_0_, *A*_l_, *A*_2_, …, *A*_*K*_, let the maternal genotype be *MG = A*_*x*_*A*_*y*_. The clutch genomic data is represented by the read-count vector *m = (m*_0_, *m*_1_, …, *m*_*K*_), where *m*_*x*_ is the observed count of allele *x* among reads from the pooled offspring sample. ***With a single sire***, the likelihood of the observed clutch read-count vector is given by:

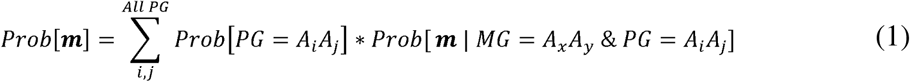

where *PG* denotes the paternal genotype and the summation is taken over all possible paternal genotypes. Assuming random mating, paternal genotype probabilities are determined from population allele frequencies:

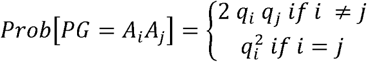

where *q_*i*_* is the population frequency of allele *i*. Conditional on maternal and paternal genotypes, the likelihood of the observed clutch read counts can be written as a sum over all possible offspring genotype-count configurations:

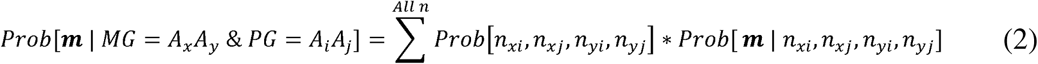

where *n*_*xi*_, *n*_*xj*_, *n*_*yi*_, and *n*_*yj*_ are the numbers of offspring with genotypes *A*_*x*_*A*_*i*_, *A*_*x*_*A*_*j*_, *A*_*y*_*A*_*i*_, and *A*_*y*_*A*_*j*_, respectively.

With ***more than one sire***, the likelihood of eq (1) is extended to all possible paternal genotype permutations across sires. Let *S* be the number of sires for a maternal family and let *w*_*i*_ be the relative contribution (fraction of offspring) of the i’th sire. The contribution weights sum to one (w_1_ + w_2_+ … + w_*S*_ — 1). For a given maternal genotype, paternal genotype combination, and contribution vector, let *Q*_*M(x)*_ be the frequency of allele *x* in the maternal genotype and *Q*_*P(x)*_ be the weighted average frequency of allele *x* among the contributing paternal genotypes. Conditional on *Q*_*M(x)*_ and *Q*_*P(x)*_, the expected read count for allele *x* in a pooled clutch sample with total read depth *M* is given by *M*[*Q*_*M(x)*_ *-I Q*_*P(x)*_]*/2*.

Observed read counts deviate from this expectation because of genetic segregation among a finite number of offspring and read sampling from the pooled clutch DNA at finite sequencing depth. With multiple sires, the number of possible paternal genotype combinations and segregation outcomes increases rapidly. To avoid summing over all possible segregation outcomes, we use an approximation based on analytical means, variances, and covariances. Let *Q*_*O(x)*_ be the realized frequency of allele *x* in the clutch pool and *n*_*d*_ be the total number of offspring in the clutch. Conditional on *Q*_*M(x)*_ and *Q*_*P(x)*_, the variance in *Q*_*O(x)*_ is:

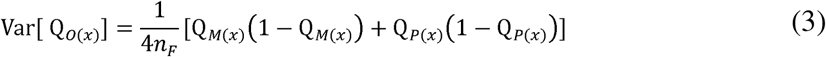

Using the variance in *Q*_*O(x)*_ from Equation (3), the variance in the read count for allele *x* is

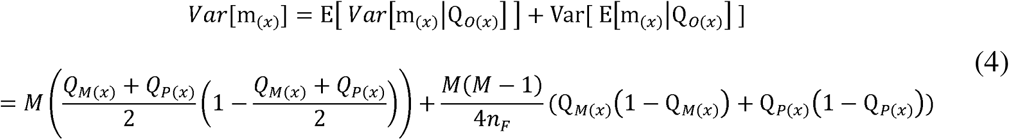

For biallelic markers, only one allele count is needed because the second allele count is determined by total read depth. Given this mean and variance, the probability of the observed allele count is approximated using a normal distribution:

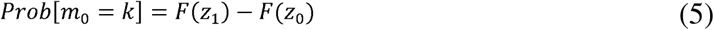

where *F*(*) is the cumulative density function of the standard normal distribution, and:

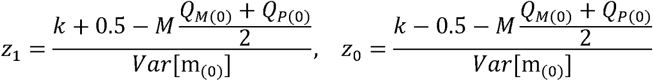

For markers with more than two alleles, we replace the univariate normal with a multivariate normal approximation. Covariances among allele counts are negative because the total read count is constrained to *M*, so an elevated count for one allele leaves less space for other alleles:

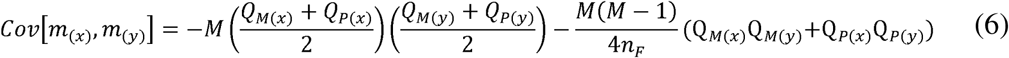

Together with the variances in Equation (4), these covariances calibrate the joint distribution of allele counts under a multivariate normal density. For four-allele markers, the joint distribution is evaluated for three allele-count classes, the fourth determined by total read depth. As with biallelic markers, the overall likelihood is calculated by averaging over all possible paternal genotype combinations.

The estimate that emerges from eqs (1)-(6) is a vector of proportions for each maternal family. For example, [0.5, 0.4, 0.1] denotes a family with three sires that contribute 50%, 40%, and 10% of the offspring, respectively. It can be informative to distill this vector into a summary statistic called the “effective number of sires” (Neff et al. 2008; Hooks and Burgess 2025). The effective number of sires is calculated as *1/(∑_*j*_f_*j*_^2^)*, where *f*_*j*_ is the fraction of offspring assigned to sire *j*, and the sum is taken over all inferred sires in the family (Bernasconi, 1993). The quantity *(∑_*j*_f_*j*_^2^)* is essentially the probability that two offspring, randomly sampled from the family, are full siblings. Effective number of sires increases with the number of sires and with evenness of their respective contributions. Assuming three sires, if all contribute equally, the effective number of sires is three, whereas if the contributions are 98%, 1%, 1%, the effective number of sires is 1.04. We use effective sire number to distill the polyandry estimate, which is a vector of proportions, into a single informative number.

### Estimation programs

The programs in our <u>github repository</u> apply the theory to obtain the likelihood of the data across all polymorphic sites for each family for every specified model of polyandry. The latter are considered as a discrete grid of possibilities for the number of sires and the fraction of offspring sired by each male. The granularity parameter determines how finely we subdivide families. For both simulations and the application, we set the granularity to 20, which implies that a specific sire can contribute 1/20, or 2/20, or 3/20 … of all offspring. Each additional sire adds one parameter to the model and so we only accept the more complicated model (say two sires with 95/5 proportions over a one sire model) if the likelihood ratio statistic is greater than 3.84 (the threshold for a chi-square distribution with 1 degree of freedom). Different proportions with the same sire number (say 50/50 versus 55/45) are models of equal complexity so the highest likelihood is selected. Full implementation details and all programs (for both real and simulated data) are available at our <u>github repository</u>.

### Simulation analyses

We used simulations to evaluate the accuracy of our estimation framework under conditions in which the true number of sires and their relative contributions were known. The simulations incorporated the two sequential sources of sampling variation described in the likelihood model: genetic segregation determining the realized genotype composition of the offspring clutch, followed by sampling of reads from the pooled clutch at finite sequencing depth. The resulting simulated data was then analyzed using the same likelihood framework applied to the empirical data.

For biallelic SNPs, the base parameter set for simulations consisted of 1000 SNPs and 200 4-allele markers with all alleles equally frequent with each marker type. There were 100 offspring per clutch and a total pooled-clutch read depth of 400 reads per marker (see program set 5 in our <u>github repository</u>). We simulated data with one-sire per families, two-sire families with male contribution proportions of 50/50, 75/25, and 90/10, and three-sire families with contribution proportions of 40/30/30 and 80/10/10. We then considered deviations from this base parameter set, variation in read depth, allele frequencies and family sizes. For each scenario, we simulated 100 maternal families, applied the routine, and then compared the polyandry estimate (sire number and paternal contribution proportions) to that specified in the simulation. We performed separate analyses using biallelic SNPs alone, four-allele markers alone, and both marker types combined.

### Drosophila field collections and clutch establishment

Wild flies were collected from plant compost material at Tableview Farm in Franklin County, Kansas, during four collection intervals in May and June of 2025. Flies were placed into collection tubes and transported to the University of Kansas, where they were anesthetized with CO□. *Drosophila melanogaster* and *D. simulans* individuals were identified visually using a dissecting microscope and separated from other flies, then species was determined by examining male offspring genitalia. Across the four collection intervals, wild mated females were individually placed into food vials and maintained for 8 days at 22°C to allow oviposition. All progeny of each female were harvested for pooled progeny sequencing.

### DNA extraction, library preparation, and sequencing

DNA Extractions were performed on field collected females and on their associated progeny pools (275 families). All extractions were done using reagents from Qiagen Puregene Tissue Kit. DNA concentration was measured using Qubit 1X dsDNA BR Assay Kit, and only samples with 7ng/ul or higher were used for library preparation. Indexed genome sequencing libraries were made using the Illumina DNA Prep Kit (Cat Num 20060059).

Sequencing libraries were prepared using an adjusted 1/10th protocol, allowing each kit to accommodate up to 10 times as many samplesThe 1/10th protocol allows for samples to be pooled before the final bead clean-up, resulting in one library plex that contains multiple samples. Female and Progeny samples were pooled into plexes of 24 to 56 samples. Samples with low DNA concentration (between 7 to 16 ng/ul) were processed into libraries using an adjusted low input 1/10th protocol. The adjusted protocol requires only 21 ng total DNA and uses 8 cycles of PCR. A subset of libraries made using this protocol were sequenced before the whole collection to verify that read counts averaged across samples as expected. The quality of all libraries was assessed using a Qubit 1X dsDNA HS Assay Kit and an Agilent TapeStation D1000 ScreenTape Assay. All libraries had fragment sizes in the range of 381bp to 568bp. Using this quality control info, we pooled libraries for sequencing. Calculations for pooling libraries were based on molecular weight, estimated reads from flow cell, and estimated genome coverage from reads. Libraries were sequenced at the Genomics Core at the University of Kansas Medical Center for sequencing using a NovaSeq X Plus 25B flow cell.

### Read preprocessing, mapping, and filtering

Raw paired-end sequencing reads were preprocessed with fastp v1.3.6 using default filtering parameters to remove adapters, trim low-quality sequence, and discard reads failing quality filters (Chen et al., 2018). Cleaned reads were then taxonomically screened using Kraken2 v2.17.1 (Wood et al., 2019) with a custom database containing the *Drosophila melanogaster* dm6 reference genome (Hoskins et al., 2015), the human reference genome (Schneider et al., 2017), and bacterial, archaeal, viral, and viroid reference sequences (O’Leary et al., 2015). Reads classified as *D. melanogaster* were extracted from the Kraken2 output using KrakenTools v1.2.1, while preserving paired-end read structure (Lu et al., 2022). The resulting filtered read pairs were used for downstream alignment and variant calling.

Filtered paired-end reads were aligned to the *Drosophila melanogaster* dm6 reference genome using the BWA-MEM algorithm v0.7.19 (Li & Durbin, 2009; Li, 2013). Alignments were converted to BAM format with *samtools view*, coordinate-sorted with *samtools sort*, and indexed with *samtools index* using SAMtools v1.24 (Danecek et al., 2021). Sample-specific read groups were added using *AddOrReplaceReadGroups* in Picard v3.3.0, with the platform specified as Illumina and the sample identifier used for both the read-group platform unit and sample name (Broad Institute, 2024). Duplicate reads were removed using Picard *MarkDuplicates* with *REMOVE_DUPLICATES — true*. To retain properly paired alignments, duplicate-filtered BAM files were name-sorted and processed with *samtools fixmate* using the *—rmu* options. Alignments were then filtered with *samtools view — f* 0*x*2 to retain reads flagged as properly paired, coordinate-sorted, and indexed. The resulting BAM files were used for variant calling.

### Variant calling, SNP filtering and marker selection

Variants were called from the filtered BAM files using bcftools mpileup with the output piped to bcftools call in BCFtools v1.24 (Danecek et al., 2021). The resulting VCF files were filtered using programs in Set 1 of our <u>github repository</u>. To avoid the inclusion of potential mapping artifacts, poorly supported sites, or uninformative variants, we excluded indels and sites segregating three bases, variants with a QUAL or mapping quality score less than 20, or with aberrant read depth. Candidate biallelic SNPs were evaluated in 10-kb windows and excluded if the Hardy–Weinberg test statistic in mothers exceeded 3.84 or minor allele frequency was less than 0.25. Because SNPs were abundant, we used the relatively restrictive minor-allele-frequency threshold to preferentially retain intermediate-frequency variants. We excluded SNPs with exaggerated frequency divergence between mothers and clutches across families and those with excessive missing data (see program Sets 1 and 2 in our <u>github repository</u>). We retained at most one SNP per 10-kb window across the genome.

After determining SNPs, we tested for congruence between the genotypes of mothers and their assigned clutches. At any SNP where the mother is homozygous, the clutch should not be segregating at frequencies significantly less than 0.5. We applied this test (probabilistically) to all SNPs as part of the SNP identification pipeline (program Set 1 in our <u>github repository</u>). To determine if the reassignments of clutches to alternative mothers were warranted, we revised the AYMM (are you my mother?) program from the BORICE mating system estimation package (Crawford, et al. 2022; https://github.com/jkkelly/BORICE.genomic.v3) to treat pooled progeny data. The program AYMM.v1.py (program Set 3 in our <u>github repository</u>) tests each clutch of a questionable mother-clutch pairing against all possible alternative clutches. For 18 of the mothers with low matching to their (initially) assigned clutch, we found very high congruence to exactly one alternative clutch. We never found high congruence to more than one alternative clutch and the reassigned clutches were always in the set that were initially identified as dubious to their respective assigned mothers. Mother-clutches reassignments were easily identified as sample swaps (e.g. mother 1_107 was reassigned to clutch 1_109, while mother 1_109 was reassigned to clutch 1_110, and mother 1_110 to clutch 1_107). Seven mother-clutch sets could not be reassigned and were eliminated. The final dataset contained 275 mother-clutch combinations.

Four-allele mini-haplotype markers were constructed from pairs of closely linked SNPs identified through the variant-calling and filtering procedures described above, using scripts adapted from Monnahan et al. (2021). Candidate SNPs were required to have a minor allele frequency of at least 0.1. So that both SNPs could be scored on the same sequencing read, only pairs occurring within a 50 bp window were considered. For each SNP pair, alleles observed on individual sequence reads were combined into one of four possible two-SNP haplotype states: RR, RA, AR, and AA. Candidate mini-haplotype markers were evaluated by comparing these haplotypes with the corresponding maternal single-SNP genotype calls and by examining haplotype patterns in mothers and progeny pools. Because each diploid mother can carry no more than two haplotypes at a two-SNP marker, loci at which three or more haplotypes were observed in any of the 275 families were excluded. Progeny pools were permitted to contain all four haplotypes but were required to contain the haplotype state or states observed in the mother. Markers were also excluded when maternal haplotypes were less abundant in progeny pools than expected under Mendelian segregation. The final four-allele mini-haplotype marker set consisted of 239 loci (programs in Set2 in our <u>github repository</u>). The likelihood model (described above) was applied to 3,053 biallelic SNPs alone, the 239 four-allele mini-haplotype markers alone, and both marker sets jointly. The 2-allele (chosen.snps1) and 4-allele (set0.4data) datasets for the 275 families that were ultimately used for analysis are available in our <u>github repository</u>.

## Results

### Simulation analyses

We first used the simulation program to illustrate how pooled clutch sequencing of biallelic SNPs can identify polyandry. Figure 2 shows the pattern for SNPs where the two alleles are equally frequent. If the mother is a homozygote, the frequency of the non-maternal allele will be distinctly trimodal with a dominant mode at zero (Fig. 2, blue), corresponding to the three possible genotypes of the sire. Each of these possibilities occurs many times across the genome yielding an accumulation of observations at zero non-maternal alleles, 25%, and 50%, with variation around the 25% and 50% modal points owing to stochastic segregation and finite read depth per locus. Most importantly, many clutches will be monomorphic for the maternal allele when the mother is a homozygote and there is a single sire (note y-axis break in Fig 2). If a second sire contributes (red trajectory in Fig 2), the frequency of monomorphic clutches is dramatically reduced (by 75% when the two alleles are equally frequent). The extreme cases of 0% and 50% non-maternal are redistributed to a greater range of intermediate possibilities. This is a key signal that enables polyandry inference with both bi-allelic and multi-allelic markers, although there is a greater range of possibilities for segregation in the latter case.

**Figure 2.**
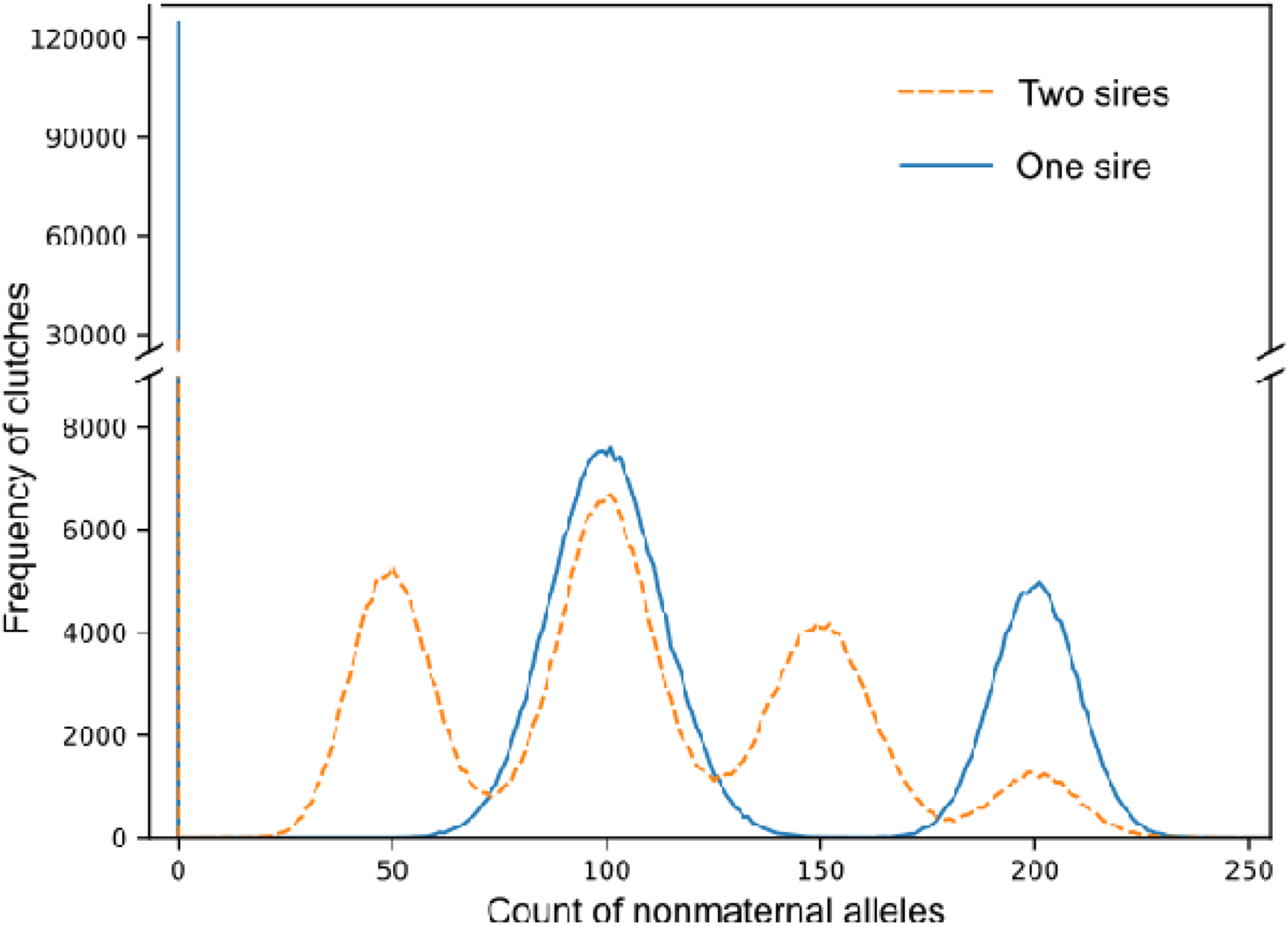
Simulation results show the difference between a single sire family (blue) and a two-sire family with equal contributions from each sire (red) for loci where the mother is homozygous. This example considers bi-allelic SNPs under the base parameter set for simulations. The count of alleles is out of 400 total per SNP.

Figure 2 simplifies the analysis for illustration. It considers only loci that are homozygous in the female, despite that maternal heterozygotes are informative and integrated into the likelihood model (eqs 1-6), and it considers biallelic SNPs with equally frequent alleles. SNPs with extreme allele frequencies are less informative. Of course, most sequencing platforms will identify a superabundance of SNPs allowing researchers to ruthlessly filter to intermediate frequency variants (our application to *Drosophila* eliminated biallelic SNPs when the minor allele was less than 25%). Finally, while Fig. 2 suggests effective discrimination between single from multi-sire families, it does not indicate efficacy of pooled-clutch sequencing for more subtle differences in sire contributions. To evaluate how finely bi- and multi- allelic markers can discriminate sire contributions, we simulated data under varying polyandry schemes.

The estimation accuracy of the likelihood model under a range of conditions is reported in Table 1. For each case, we compared the error rate when using only bi-allelic markers, only 4-allele markers, and both combined. Consistent with Figure 2, the method always discriminate single sire families from multiple sire families (no errors across families for any parameter set). Estimation was generally correct for the base parameter set (section A of Table 1) except for the case of three sire families with 40/30/30 proportions. Here, only about a third of family specific estimates exactly match the truth, with two thirds deviating by a single step. The most frequently observed incorrect estimate was 35/35/30, which yielded an absolute error score of 0.033. The average absolute error rate was near 0.02 for all three marker sets, with about a third of the families having absolute error 0.0, and the rest having a one-step error.

**Table 1.** For each of 5 parameter sets (A-E), simulation results from 100 families are reported for each of six different polyandry models (one sire, two sires, or three sires with differing proportions in the latter cases). In each cell, the first number is the percentage of estimates exactly matching the specified proportions. The second is the average (absolute) error in the estimated sire contributions.

|  | Polyandry model | Model parameters | Only SNPs | Only 4-allele markers | Both |
| --- | --- | --- | --- | --- | --- |
| (A) | One Sire | Base | 100% / 0.0 | 100% / 0.0 | 100% / 0.0 |
|  | Two Sires (50/50) | Base | 100% / 0.0 | 100% / 0.0 | 100% / 0.0 |
|  | Two Sires (75/25) | Base | 100% / 0.0 | 100% / 0.0 | 100% / 0.0 |
|  | Two Sires (90/10) | Base | 100% / 0.0 | 100% / 0.0 | 100% / 0.0 |
|  | Three Sires (40/30/30) | Base | 36% / 0.021 | 40% / 0.020 | 37% / 0.021 |
|  | Three Sires (80/10/10) | Base | 99% / 0.0003 | 100% / 0.0 | 100% / 0.0 |
| (B) | One Sire | Read depth = 0.5x | 100% / 0.0 | 100% / 0.0 | 100% / 0.0 |
|  | Two Sires (50/50) | Read depth = 0.5x | 88% / 0.007 | 79% / 0.011 | 94% / 0.003 |
|  | Two Sires (75/25) | Read depth = 0.5x | 89% / 0.006 | 91% / 0.005 | 99% / 0.0005 |
|  | Two Sires (90/10) | Read depth = 0.5x | 100% / 0.0 | 100% / 0.0 | 99% / 0.0003 |
|  | Three Sires (40/30/30) | Read depth = 0.5x | 14% / 0.031 | 33% / 0.023 | 30% / 0.024 |
|  | Three Sires (80/10/10) | Read depth = 0.5x | 88% / 0.004 | 88% / 0.004 | 94% / 0.002 |
| (C) | One Sire | Read depth = 0.5x, nf = 400 | 100% / 0.0 | 100% / 0.0 | 100% / 0.0 |
|  | Two Sires (50/50) | Read depth = 0.5x, nf = 400 | 100% / 0.0 | 100% / 0.0 | 100% / 0.0 |
|  | Two Sires (75/25) | Read depth = 0.5x, nf = 400 | 100% / 0.0 | 100% / 0.0 | 100% / 0.0 |
|  | Two Sires (90/10) | Read depth = 0.5x, nf = 400 | 100% / 0.0 | 100% / 0.0 | 100% / 0.0 |
|  | Three Sires (40/30/30) | Read depth = 0.5x, nf = 400 | 82% / 0.006 | 86% / 0.005 | 94% / 0.002 |
|  | Three Sires (80/10/10) | Read depth = 0.5x, nf = 400 | 99% / 0.00033 | 100% / 0.0 | 100% / 0.0 |
| (D) | One Sire | Uneven allele frequencies | 100% / 0.0 | 100% / 0.0 | 100% / 0.0 |
|  | Two Sires (50/50) | Uneven allele frequencies | 100% / 0.0 | 99% / 0.0003 | 100% / 0.0 |
|  | Two Sires (75/25) | Uneven allele frequencies | 100% / 0.0 | 100% / 0.0 | 100% / 0.0 |
|  | Two Sires (90/10) | Uneven allele frequencies | 100% / 0.0 | 100% / 0.0 | 100% / 0.0 |
|  | Three Sires (40/30/30) | Uneven allele frequencies | 35% / 0.022 | 28% / 0.024 | 26% / 0.025 |
|  | Three Sires (80/10/10) | Uneven allele frequencies | 100% / 0.0 | 100% / 0.0 | 100% / 0.0 |
| (E) | One Sire | fam size = 20 | 100% / 0.0 | 100% / 0.0 | 100% / 0.0 |
|  | Two Sires (50/50) | fam size = 20 | 87% / 0.0095 | 84% / 0.0125 | 94% / 0.003 |
|  | Two Sires (75/25) | fam size = 20 | 40% / 0.033 | 69% / 0.0155 | 61% / 0.0195 |
|  | Two Sires (90/10) | fam size = 20 | 96% / 0.002 | 96% / 0.002 | 100% / 0.0 |
|  | Three Sires (40/30/30) | fam size = 20 | 2% / 0.033 | 8% / 0.0317 | 1% / 0.033 |
|  | Three Sires (80/10/10) | fam size = 20 | 77% / 0.011 | 60% / 0.0183 | 88% / 0.005 |

We next considered reducing the sequencing depth from two reads per chromosome per clutch (400x over 100 diploid offspring) to 0.5x per chromosome (section B in Table 1). Predictably, this increased error rates, but only slightly in quantitative terms. For example, with two sires and 50/50 proportions, combined markers were perfect for the base parameter set but not for 6 of 100 families with 0.5x coverage. However, all 6 families were estimated as 55/45 and so the effective number of sires was underestimated by only 1% in these families. Parameter Set C held the coverage per chromosome to 0.5x (as in section B) but increased family size (400 instead of 100) so that the total sequence depth per clutch matched the base parameter set. The result is improved accuracy, particularly for the most error prone case of three sires with 40/30/30, suggesting that increased clutch size may be more beneficial than increased sequencing coverage per individual within a clutch Next, we considered skewed allele frequencies. For bi-allelic markers, we simulated data with 75/25 allele frequencies for bi-allelic markers and 70/10/10/10 for 4-allele markers (instead of 50/50 and 25/25/25/25 as with the base parameter set). We found that this level of asymmetry in allele frequencies had a negligible effect on error rates (compare sections D to A in table 1). Of course, extreme allele frequency skew (e.g. 99/1) will make markers less informative because the minor allele will be absent from most families regardless of mating the system. However, the scale of genomic data means that filtering out extremely skewed markers should still leave a saturating abundance of markers. Finally, we considered smaller families (20 instead of 100, section E of Table 1), which yielded the highest error rates. Error was not large in absolute terms (almost never one step away from the truth), but imperfect inference was more frequent for various 2-sire scenarios.

Within a polyandry scenario (a single row of Table 1), estimation accuracy fluctuated between 1000 SNPs considered alone, 200 4-allele makers considered alone, and the combined marker set. Some of this variation was stochastic (only 100 families scored per case). Across all 30 cases, the mean absolute error for SNPs (0.0062) was slightly greater than for 4-allele markers (0.0057), emphasizing the informativeness of multi-allelic markers for paternity inference. Using the two sets in combination was considerably better than either alone, evidenced by a ∼25% reduction in the mean absolute error (0.0046). Notably, these simulations used fewer loci than were available in the empirical application below. These numbers should be readily attainable with genome-wide genotyping in most species.

### Empirical estimates of polyandry

We estimated polyandry for 275 wild-caught mother-clutch families, first using the data from biallelic SNPs and then using the four-allele mini-haplotypes. Estimates based on the 3,053 biallelic SNPs were highly concordant with estimates based on the 239 four-allele mini-haplotype markers. Sire number estimates were identical for 250 of 275 families (91%). When estimates differed (25 families), the differences involved minor changes in inferred paternal contribution rather than large differences in effective number of sires. For example, in family 1_31, the four-allele markers supported a single-sire model, whereas the biallelic SNPs supported a two-sire model in which the primary sire contributed 95% of the progeny. Mean effective number of sires was 1.16 for the biallelic SNP analysis and 1.14 for the four-allele marker analysis. The mean absolute difference in effective number of sires between marker sets was 0.033, indicating that the two marker sets produced nearly identical estimates of effective sire number per family (Fig. 3A). The full detail for family specific model fits is reported for markers combined in Supplemental Tables S1, and for each marker set separately as Supplemental Tables S2,S3).

**Figure 3.**
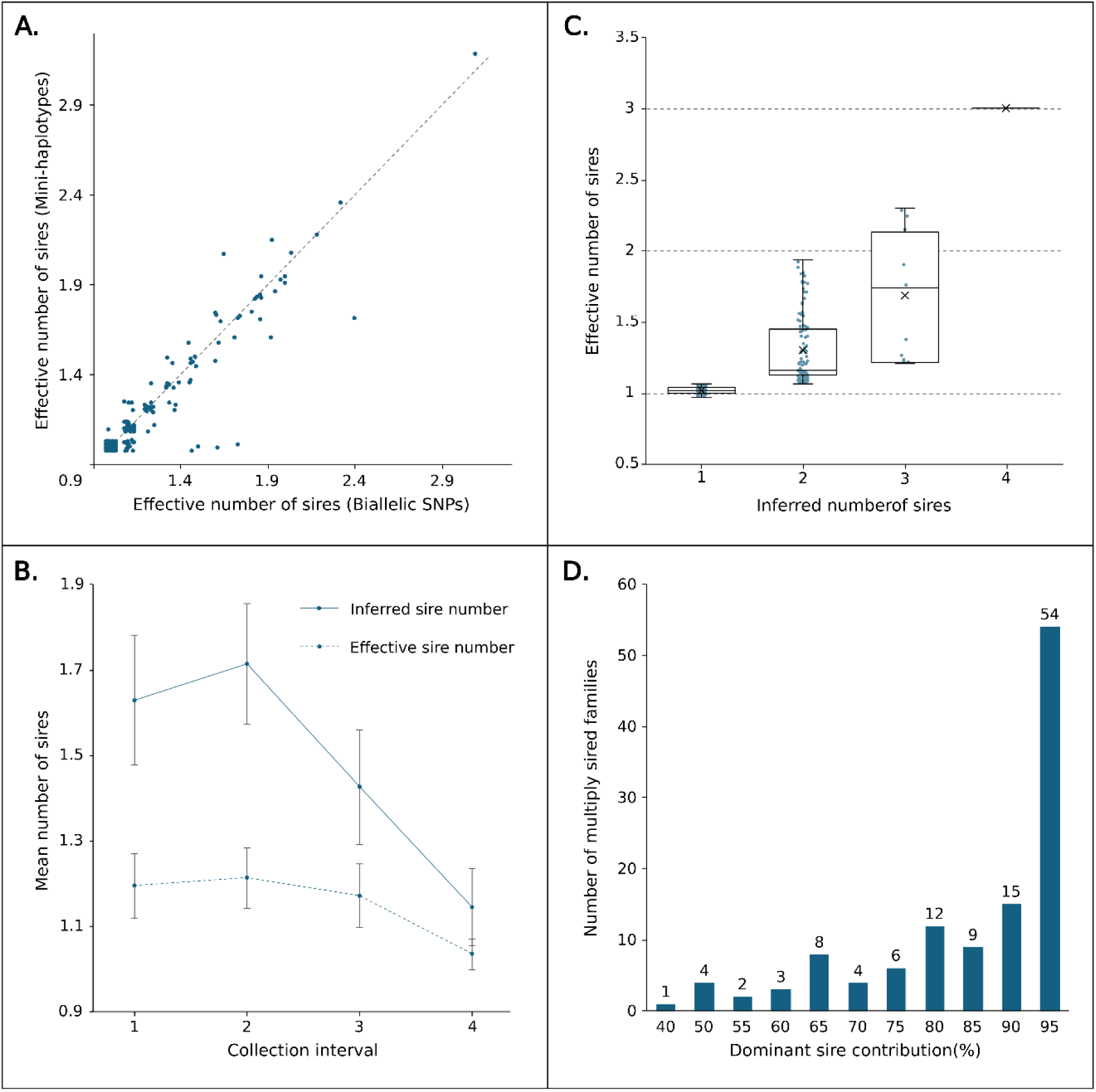
Empirical estimates of multiple paternity among 275 wild-caught *Drosophila melanogaster* maternal families. A) Concordance between effective number of sires estimated independently from 3,053 biallelic SNPs and 239 four-allele mini-haplotype markers. Each point represents one family; points were slightly offset to improve visibility. B) Mean inferred sire number (solid line) and effective number of sires (dashed line) across the four collection intervals. Error bars indicate 95% confidence intervals. Collection interval had a significant effect on both inferred sire number (P < 0.001) and effective number of sires (P = 0.002). C) Distribution of effective number of sires among families assigned one to four inferred sires. Points represent individual families and were slightly offset in both dimensions to improve visibility. D) Distribution of the inferred contribution of the dominant sire among the 118 families showing evidence of multiple paternity. Bar labels indicate the number of families in each paternal-contribution class.

We then combined 2-allele and 4-allele markers to obtain the final empirical estimate for each family. Inferred sire number ranged from one to four, but estimates were concentrated at one or two sires: 157 families had one inferred sire, 106 had two sires, 11 had three sires, and one had four sires (Fig. 3C). Thus, 118 of 275 families showed evidence of multiple paternity. Among multiply sired families, the inferred dominant sire contributed an average of 84.6% of offspring, indicating that multiple paternity was often highly skewed (Fig. 3D). Across all families, the mean effective number of sires was 1.16, with a range of 1 to 3.2. Effective number of sires was concentrated near one: 157 families had an effective number of sires of exactly 1, 113 had values >1 to 2, and 5 had values >2. Family size, which ranged from 8 to 137 individuals per progeny pool, was not correlated with inferred sire number (*r = 0.105, P = 0.81*) or effective number of sires (*r = —0.017, P = 0.77*), indicating that higher sire-number estimates were not simply a consequence of larger families.

Polyandry estimates varied across collection intervals even though these collections were only a few days apart. Mean inferred sire number was 1.63 in collection 1, 1.71 in collection 2, 1.43 in collection 3, and 1.15 in collection 4. Mean effective number of sires were 1.20, 1.21, 1.17, and 1.04, respectively. Collection interval had a significant effect on both inferred sire number (*P <! 0.001*) and effective number of sires (*P = 0.002*). Mean estimates were highest in the first two collection intervals and lower in later collections, consistent with temporal variation in realized multiple paternity (Fig. 3B).

## Discussion

Multiple mating by females (polyandry) can increase fecundity and genetic diversity within a clutch. The level of polyandry has critical consequences for processes such as kin selection and sperm competition. Multiple mating partitions a maternal family into multiple paternal sub-families, each composed of full siblings. Our premise is that one can detect this internal structure without genotyping individual offspring. Pooled sequencing of maternal families can effectively delineate the number and relative contribution of sires by leveraging the subtle signature of Mendelian segregation across hundreds to thousands of loci. This enables the quantitative characterization of polyandry across a diverse range of species with large maternal families where individual progeny genotyping is inpractical.

Simulations show that polyandry, measured in terms of sire number and relative paternal contribution, can be accurately estimated using genomic data from individual mothers and their corresponding pooled progeny (Table 1). Accuracy is especially high for one- and two-sire families, even when inference is based only on biallelic SNPs. This result is especially encouraging given that simulations used only 1000 SNPs, a relatively small marker set relative to the tens of thousands of markers commonly generated by genome-wide sequencing approaches (Davey et al., 2011). Estimation accuracy was marginally lower with fewer offspring per clutch (Table 1E). In part, this simply reflects a lower volume of data than is available when larger clutches (all else equal). However, smaller families also strain one aspect of the theory. The model relies on a continuous probability density (the multi-normal) to approximate likelihood for discrete data (read counts sampled from a finite number of offspring). The assumption is critical for obtaining the likelihood without summing over the enormous number parental genotype and segregation possibilities. Discrepancies between the approximate and actual likelihoods are expected with smaller families but we have found them to be quantitatively small in simulations using families of 10-20 in size. In organisms that routinely have fewer than 10 offspring per clutch, individual progeny genotyping may be preferrable anyway.

In our application to *Drosophila*, estimates from the biallelic SNPs and four-allele mini-haplotypes were highly concordant: sire-number estimates were identical for 250 of 275 families, and the mean absolute difference between marker-set estimates was only 0.033 effective sires per family (Fig 3A). This agreement provides an important check on the stability of the inference. The success of the mini-haplotype analysis also shows that short read sequencing can provide multi-allelic information analogous to traditional microsatellite markers from high-throughput short-read sequencing (Baetscher et al., 2017; Flanagan & Jones, 2019). We exploit the scale of genomic data for marker selection. For example, our initial scan identified over 370,000 SNPs that were within 50bp of another SNP. This enabled very stringent selection of 4-allele markers based on mother-offspring consistency, data completeness, and allele frequencies (all four haplotypes must be abundant in the experiment).

Estimates of polyandry from the Kansas population are broadly consistent with previous studies of multiple paternity in wild *Drosophila melanogaster* (Fig 3). Using allozymes, Ochando et al. (1996) found evidence for multiple paternity in two *Drosophila melanogaster* populations, with at least two sires detected in ∼50% of maternal families. Harshman and Clark (1998) estimated an average of approximately 1.8 sires per brood in field-caught females, with a maximum inferred sire number of four. Good et al. (2006), using highly polymorphic microsatellite loci, found higher levels of multiple paternity in another fruit fly, *Drosophila mojavensis*. Our results are therefore broadly consistent with earlier evidence that multiple paternity is common in wild *Drosophila* populations. As in previous studies, we found that up to four sires can contribute to a single clutch, and that paternal contributions are often highly uneven. The pooled sequencing of full clutches enables larger studies (more maternal families) while avoiding the potential biases associated with progeny subsampling (Quennessen et al., 2025).

*Drosophila* families exhibit a strong skew in paternal contributions (Birkhead & Pizzari, 2002; Kvarnemo & Simmons, 2013). Among multiply sired families, the inferred dominant sire contributed an average of 84.6% of offspring, indicating that multiple paternity was often highly unequal (Fig 3C & Fig 3D). Because offspring are genotyped after fertilization, we are estimating realized paternity after post-copulatory processes have acted. Highly skewed families, such as those with inferred contributions near 95/5, could therefore reflect sperm competition, sperm displacement, cryptic female choice, differential fertilization success, or early offspring viability differences (Birkhead & Pizzari, 2002; Simmons, 2005). The similarity between our observed paternal skew and previous estimates of sperm displacement is consistent with the possibility that the dominant sire often represents the most recent or most successful mate, but mating order was not observed directly, so this interpretation remains tentative (Harshman & Clark, 1998; Clark & Begun, 1998). Polyandry estimates also varied across collection intervals, with higher mean sire-number and effective-sire-number estimates in the first two collection intervals than in later collections (Fig 3B). This pattern suggests temporal variation in realized paternity (Behrman et al., 2015), but it may also be related to variation in collection/processing time.

The motivation for pooled progeny sequencing is to accommodate species with large families. In organisms that produce relatively few offspring, direct genotyping of all progenies is both feasible and effective. In species that produce tens, hundreds, or thousands of offspring, individual genotyping becomes costly, and subsampling progeny can miss minor sires or distort estimates of paternal contribution (Eriksson et al., 2009; Quennessen et al., 2025). Pooled progeny sequencing is useful for species with high reproductive output and where the distribution of paternal contributions may be an important component of fitness, sexual selection, and post-copulatory competition (Birkhead & Pizzari, 2002; Kvarnemo & Simmons, 2013). In our empirical data, family size was not significantly correlated with inferred sire number or effective number of sires, indicating that higher sire-number estimates were not simply a consequence of larger progeny pools.

A disadvantage of pooled sequencing is the inevitable loss of information. Our method can determine that a maternal family is derived from two sires, but not which individual offspring are derived from each sire. The partitioning of maternal families into sub-families of full siblings, which can be determined with individual progeny genotyping (e.g. Colicchio et al., 2019; Kerbs et al., 2020), is impossible with pooled sequencing. Pooling can also introduce errors. Our simulations indicate accurate estimation with only moderate numbers of loci (Table 1) but we assumed that individual progeny contribute equally to the DNA pool. Variation in offspring size, extraction efficiency, or library preparation could shift pooled allele frequencies and affect estimates of paternal contribution. Finally, because clutches are collected from wild-caught females, the number and order of matings before collection are unknown, as is the relationship between mating order and inferred sire contribution. The method estimates only realized paternity and not the complete mating history.

## Acknowledgements

We acknowledge grant NSF IOS-2421689 and NIH grant P20-GM103418 for supporting these studies. We particularly thank Robert Hickerson for access to his farm for yearly collections.

## Notes

https://github.com/jkkelly/Polyandry_by_pooled_families

### Competing Interest Statement

The authors have declared no competing interest.

