## Supplemental Tables for "Pooled progeny sequencing effectively estimates polyandry in species with large families"

**Supplemental Table S1. The estimated polyandry for each family is reported in terms of proportion to each sire (w1 to w4). We also report the collection for each family, the number of offspring per clutch and the effective sire number (which is just a function of the w values).**

| Collection | Family | Number offspring | No Sires | Proportions | | | | Effective number sires |
| --- | --- | --- | --- | --- | --- | --- | --- | --- |
|  |  |  |  | w1 | w2 | w3 | w4 |  |
| 1 | 1_101 | 16 | 2 | 0.65 | 0.35 |  |  | 1.83 |
| 1 | 1_102 | 40 | 2 | 0.85 | 0.15 |  |  | 1.34 |
| 1 | 1_105 | 11 | 1 | 1 |  |  |  | 1.00 |
| 1 | 1_108 | 21 | 1 | 1 |  |  |  | 1.00 |
| 1 | 1_11 | 20 | 2 | 0.95 | 0.05 |  |  | 1.10 |
| 1 | 1_111 | 25 | 2 | 0.75 | 0.25 |  |  | 1.60 |
| 1 | 1_116 | 79 | 2 | 0.95 | 0.05 |  |  | 1.10 |
| 1 | 1_12 | 104 | 3 | 0.9 | 0.05 | 0.05 |  | 1.23 |
| 1 | 1_13 | 36 | 1 | 1 |  |  |  | 1.00 |
| 1 | 1_19 | 41 | 2 | 0.95 | 0.05 |  |  | 1.10 |
| 1 | 1_20 | 24 | 1 | 1 |  |  |  | 1.00 |
| 1 | 1_24 | 28 | 1 | 1 |  |  |  | 1.00 |
| 1 | 1_25 | 34 | 1 | 1 |  |  |  | 1.00 |
| 1 | 1_26 | 87 | 2 | 0.95 | 0.05 |  |  | 1.10 |
| 1 | 1_27 | 39 | 1 | 1 |  |  |  | 1.00 |
| 1 | 1_28 | 59 | 2 | 0.95 | 0.05 |  |  | 1.10 |
| 1 | 1_31 | 46 | 2 | 0.95 | 0.05 |  |  | 1.10 |
| 1 | 1_33 | 71 | 2 | 0.9 | 0.1 |  |  | 1.22 |
| 1 | 1_38 | 39 | 1 | 1 |  |  |  | 1.00 |
| 1 | 1_39 | 62 | 2 | 0.95 | 0.05 |  |  | 1.10 |
| 1 | 1_4 | 11 | 2 | 0.8 | 0.2 |  |  | 1.47 |
| 1 | 1_40 | 72 | 2 | 0.95 | 0.05 |  |  | 1.10 |
| 1 | 1_42 | 53 | 1 | 1 |  |  |  | 1.00 |
| 1 | 1_45 | 57 | 2 | 0.95 | 0.05 |  |  | 1.10 |
| 1 | 1_46 | 43 | 2 | 0.5 | 0.5 |  |  | 2.00 |
| 1 | 1_47 | 58 | 2 | 0.9 | 0.1 |  |  | 1.22 |
| 1 | 1_5 | 32 | 1 | 1 |  |  |  | 1.00 |
| 1 | 1_56 | 17 | 1 | 1 |  |  |  | 1.00 |
| 1 | 1_58 | 38 | 2 | 0.95 | 0.05 |  |  | 1.10 |
| 1 | 1_61 | 44 | 2 | 0.85 | 0.15 |  |  | 1.34 |
| 1 | 1_63 | 77 | 1 | 1 |  |  |  | 1.00 |
| 1 | 1_64 | 29 | 2 | 0.95 | 0.05 |  |  | 1.10 |
| 1 | 1_65 | 22 | 2 | 0.8 | 0.2 |  |  | 1.47 |
| 1 | 1_67 | 23 | 1 | 1 |  |  |  | 1.00 |
| 1 | 1_69 | 49 | 2 | 0.95 | 0.05 |  |  | 1.10 |
| 1 | 1_70 | 21 | 2 | 0.9 | 0.1 |  |  | 1.22 |
| 1 | 1_71 | 30 | 2 | 0.55 | 0.45 |  |  | 1.98 |
| 1 | 1_72 | 83 | 3 | 0.65 | 0.3 | 0.05 |  | 1.94 |
| 1 | 1_73 | 35 | 2 | 0.75 | 0.25 |  |  | 1.60 |
| 1 | 1_74 | 59 | 2 | 0.85 | 0.15 |  |  | 1.34 |
| 1 | 1_75 | 39 | 2 | 0.9 | 0.1 |  |  | 1.22 |
| 1 | 1_76 | 47 | 2 | 0.65 | 0.35 |  |  | 1.83 |
| 1 | 1_79 | 64 | 1 | 1 |  |  |  | 1.00 |
| 1 | 1_8 | 32 | 1 | 1 |  |  |  | 1.00 |
| 1 | 1_80 | 43 | 1 | 1 |  |  |  | 1.00 |
| 1 | 1_83 | 44 | 1 | 1 |  |  |  | 1.00 |
| 1 | 1_86 | 38 | 2 | 0.9 | 0.1 |  |  | 1.22 |
| 1 | 1_88 | 29 | 1 | 1 |  |  |  | 1.00 |
| 1 | 1_89 | 21 | 1 | 1 |  |  |  | 1.00 |
| 1 | 1_90 | 64 | 2 | 0.95 | 0.05 |  |  | 1.10 |
| 1 | 1_91 | 20 | 1 | 1 |  |  |  | 1.00 |
| 1 | 1_93 | 68 | 1 | 1 |  |  |  | 1.00 |
| 1 | 1_94 | 31 | 2 | 0.95 | 0.05 |  |  | 1.10 |
| 1 | 1_97 | 67 | 1 | 1 |  |  |  | 1.00 |
| 2 | 2_10 | 82 | 3 | 0.9 | 0.05 | 0.05 |  | 1.23 |
| 2 | 2_100 | 92 | 2 | 0.8 | 0.2 |  |  | 1.47 |
| 2 | 2_101 | 41 | 1 | 1 |  |  |  | 1.00 |
| 2 | 2_102 | 62 | 2 | 0.9 | 0.1 |  |  | 1.22 |
| 2 | 2_103 | 33 | 1 | 1 |  |  |  | 1.00 |
| 2 | 2_105 | 49 | 2 | 0.85 | 0.15 |  |  | 1.34 |
| 2 | 2_108 | 61 | 1 | 1 |  |  |  | 1.00 |
| 2 | 2_12 | 66 | 1 | 1 |  |  |  | 1.00 |
| 2 | 2_122 | 58 | 2 | 0.95 | 0.05 |  |  | 1.10 |
| 2 | 2_123 | 30 | 2 | 0.8 | 0.2 |  |  | 1.47 |
| 2 | 2_125 | 24 | 1 | 1 |  |  |  | 1.00 |
| 2 | 2_127 | 26 | 3 | 0.5 | 0.4 | 0.1 |  | 2.38 |
| 2 | 2_132 | 42 | 2 | 0.95 | 0.05 |  |  | 1.10 |
| 2 | 2_136 | 43 | 2 | 0.75 | 0.25 |  |  | 1.60 |
| 2 | 2_137 | 83 | 1 | 1 |  |  |  | 1.00 |
| 2 | 2_14 | 44 | 2 | 0.85 | 0.15 |  |  | 1.34 |
| 2 | 2_16 | 65 | 1 | 1 |  |  |  | 1.00 |
| 2 | 2_2 | 64 | 3 | 0.9 | 0.05 | 0.05 |  | 1.23 |
| 2 | 2_25 | 81 | 1 | 1 |  |  |  | 1.00 |
| 2 | 2_26 | 59 | 2 | 0.95 | 0.05 |  |  | 1.10 |
| 2 | 2_27 | 78 | 2 | 0.65 | 0.35 |  |  | 1.83 |
| 2 | 2_28 | 71 | 1 | 1 |  |  |  | 1.00 |
| 2 | 2_29 | 43 | 2 | 0.75 | 0.25 |  |  | 1.60 |
| 2 | 2_3 | 72 | 2 | 0.8 | 0.2 |  |  | 1.47 |
| 2 | 2_30 | 49 | 2 | 0.95 | 0.05 |  |  | 1.10 |
| 2 | 2_32 | 58 | 2 | 0.95 | 0.05 |  |  | 1.10 |
| 2 | 2_33 | 51 | 2 | 0.95 | 0.05 |  |  | 1.10 |
| 2 | 2_34 | 54 | 2 | 0.95 | 0.05 |  |  | 1.10 |
| 2 | 2_35 | 86 | 3 | 0.5 | 0.45 | 0.05 |  | 2.20 |
| 2 | 2_36 | 63 | 2 | 0.95 | 0.05 |  |  | 1.10 |
| 2 | 2_38 | 58 | 2 | 0.95 | 0.05 |  |  | 1.10 |
| 2 | 2_39 | 61 | 1 | 1 |  |  |  | 1.00 |
| 2 | 2_42 | 39 | 1 | 1 |  |  |  | 1.00 |
| 2 | 2_43 | 80 | 2 | 0.95 | 0.05 |  |  | 1.10 |
| 2 | 2_45 | 84 | 2 | 0.95 | 0.05 |  |  | 1.10 |
| 2 | 2_49 | 72 | 1 | 1 |  |  |  | 1.00 |
| 2 | 2_50 | 74 | 1 | 1 |  |  |  | 1.00 |
| 2 | 2_51 | 32 | 1 | 1 |  |  |  | 1.00 |
| 2 | 2_52 | 107 | 2 | 0.95 | 0.05 |  |  | 1.10 |
| 2 | 2_53 | 75 | 2 | 0.7 | 0.3 |  |  | 1.72 |
| 2 | 2_54 | 41 | 1 | 1 |  |  |  | 1.00 |
| 2 | 2_55 | 66 | 2 | 0.95 | 0.05 |  |  | 1.10 |
| 2 | 2_57 | 115 | 2 | 0.9 | 0.1 |  |  | 1.22 |
| 2 | 2_58 | 86 | 2 | 0.95 | 0.05 |  |  | 1.10 |
| 2 | 2_59 | 76 | 2 | 0.95 | 0.05 |  |  | 1.10 |
| 2 | 2_6 | 38 | 1 | 1 |  |  |  | 1.00 |
| 2 | 2_64 | 90 | 1 | 1 |  |  |  | 1.00 |
| 2 | 2_66 | 90 | 1 | 1 |  |  |  | 1.00 |
| 2 | 2_68 | 59 | 2 | 0.9 | 0.1 |  |  | 1.22 |
| 2 | 2_69 | 52 | 2 | 0.95 | 0.05 |  |  | 1.10 |
| 2 | 2_7 | 21 | 2 | 0.7 | 0.3 |  |  | 1.72 |
| 2 | 2_70 | 31 | 2 | 0.95 | 0.05 |  |  | 1.10 |
| 2 | 2_71 | 78 | 2 | 0.95 | 0.05 |  |  | 1.10 |
| 2 | 2_72 | 34 | 1 | 1 |  |  |  | 1.00 |
| 2 | 2_73 | 91 | 1 | 1 |  |  |  | 1.00 |
| 2 | 2_75 | 74 | 2 | 0.95 | 0.05 |  |  | 1.10 |
| 2 | 2_76 | 93 | 2 | 0.8 | 0.2 |  |  | 1.47 |
| 2 | 2_78 | 30 | 2 | 0.6 | 0.4 |  |  | 1.92 |
| 2 | 2_79 | 103 | 3 | 0.9 | 0.05 | 0.05 |  | 1.23 |
| 2 | 2_80 | 91 | 1 | 1 |  |  |  | 1.00 |
| 2 | 2_83 | 27 | 2 | 0.8 | 0.2 |  |  | 1.47 |
| 2 | 2_84 | 41 | 1 | 1 |  |  |  | 1.00 |
| 2 | 2_87 | 48 | 2 | 0.7 | 0.3 |  |  | 1.72 |
| 2 | 2_88 | 35 | 2 | 0.85 | 0.15 |  |  | 1.34 |
| 2 | 2_92 | 108 | 1 | 1 |  |  |  | 1.00 |
| 2 | 2_93 | 37 | 1 | 1 |  |  |  | 1.00 |
| 2 | 2_94 | 38 | 2 | 0.85 | 0.15 |  |  | 1.34 |
| 2 | 2_95 | 132 | 1 | 1 |  |  |  | 1.00 |
| 2 | 2_96 | 110 | 2 | 0.95 | 0.05 |  |  | 1.10 |
| 2 | 2_98 | 41 | 2 | 0.95 | 0.05 |  |  | 1.10 |
| 3 | 3_1 | 65 | 1 | 1 |  |  |  | 1.00 |
| 3 | 3_102 | 65 | 2 | 0.8 | 0.2 |  |  | 1.47 |
| 3 | 3_107 | 23 | 1 | 1 |  |  |  | 1.00 |
| 3 | 3_108 | 73 | 2 | 0.95 | 0.05 |  |  | 1.10 |
| 3 | 3_111 | 66 | 1 | 1 |  |  |  | 1.00 |
| 3 | 3_113 | 21 | 3 | 0.85 | 0.1 | 0.05 |  | 1.36 |
| 3 | 3_115 | 36 | 2 | 0.9 | 0.1 |  |  | 1.22 |
| 3 | 3_116 | 68 | 1 | 1 |  |  |  | 1.00 |
| 3 | 3_117 | 37 | 1 | 1 |  |  |  | 1.00 |
| 3 | 3_118 | 71 | 1 | 1 |  |  |  | 1.00 |
| 3 | 3_119 | 48 | 1 | 1 |  |  |  | 1.00 |
| 3 | 3_121 | 57 | 2 | 0.75 | 0.25 |  |  | 1.60 |
| 3 | 3_122 | 74 | 1 | 1 |  |  |  | 1.00 |
| 3 | 3_126 | 22 | 1 | 1 |  |  |  | 1.00 |
| 3 | 3_128 | 41 | 1 | 1 |  |  |  | 1.00 |
| 3 | 3_129 | 28 | 2 | 0.8 | 0.2 |  |  | 1.47 |
| 3 | 3_130 | 26 | 1 | 1 |  |  |  | 1.00 |
| 3 | 3_131 | 82 | 1 | 1 |  |  |  | 1.00 |
| 3 | 3_132 | 30 | 1 | 1 |  |  |  | 1.00 |
| 3 | 3_133 | 34 | 1 | 1 |  |  |  | 1.00 |
| 3 | 3_134 | 14 | 1 | 1 |  |  |  | 1.00 |
| 3 | 3_135 | 55 | 1 | 1 |  |  |  | 1.00 |
| 3 | 3_136 | 87 | 1 | 1 |  |  |  | 1.00 |
| 3 | 3_140 | 30 | 2 | 0.95 | 0.05 |  |  | 1.10 |
| 3 | 3_142 | 58 | 2 | 0.95 | 0.05 |  |  | 1.10 |
| 3 | 3_144 | 51 | 1 | 1 |  |  |  | 1.00 |
| 3 | 3_147 | 20 | 1 | 1 |  |  |  | 1.00 |
| 3 | 3_148 | 97 | 3 | 0.55 | 0.35 | 0.1 |  | 2.30 |
| 3 | 3_149 | 75 | 1 | 1 |  |  |  | 1.00 |
| 3 | 3_152 | 29 | 2 | 0.8 | 0.2 |  |  | 1.47 |
| 3 | 3_154 | 42 | 1 | 1 |  |  |  | 1.00 |
| 3 | 3_155 | 42 | 1 | 1 |  |  |  | 1.00 |
| 3 | 3_156 | 57 | 2 | 0.95 | 0.05 |  |  | 1.10 |
| 3 | 3_157 | 19 | 2 | 0.95 | 0.05 |  |  | 1.10 |
| 3 | 3_16 | 100 | 1 | 1 |  |  |  | 1.00 |
| 3 | 3_161 | 41 | 1 | 1 |  |  |  | 1.00 |
| 3 | 3_163 | 75 | 2 | 0.75 | 0.25 |  |  | 1.60 |
| 3 | 3_165 | 41 | 1 | 1 |  |  |  | 1.00 |
| 3 | 3_166 | 36 | 2 | 0.95 | 0.05 |  |  | 1.10 |
| 3 | 3_167 | 81 | 1 | 1 |  |  |  | 1.00 |
| 3 | 3_168 | 67 | 1 | 1 |  |  |  | 1.00 |
| 3 | 3_17 | 68 | 2 | 0.95 | 0.05 |  |  | 1.10 |
| 3 | 3_171 | 80 | 1 | 1 |  |  |  | 1.00 |
| 3 | 3_172 | 104 | 2 | 0.95 | 0.05 |  |  | 1.10 |
| 3 | 3_173 | 8 | 1 | 1 |  |  |  | 1.00 |
| 3 | 3_174 | 35 | 1 | 1 |  |  |  | 1.00 |
| 3 | 3_175 | 13 | 1 | 1 |  |  |  | 1.00 |
| 3 | 3_18 | 43 | 2 | 0.8 | 0.2 |  |  | 1.47 |
| 3 | 3_181 | 28 | 1 | 1 |  |  |  | 1.00 |
| 3 | 3_182 | 47 | 1 | 1 |  |  |  | 1.00 |
| 3 | 3_184 | 44 | 1 | 1 |  |  |  | 1.00 |
| 3 | 3_20 | 73 | 1 | 1 |  |  |  | 1.00 |
| 3 | 3_21 | 54 | 1 | 1 |  |  |  | 1.00 |
| 3 | 3_24 | 55 | 1 | 1 |  |  |  | 1.00 |
| 3 | 3_25 | 38 | 1 | 1 |  |  |  | 1.00 |
| 3 | 3_26 | 30 | 2 | 0.85 | 0.15 |  |  | 1.34 |
| 3 | 3_3 | 38 | 1 | 1 |  |  |  | 1.00 |
| 3 | 3_31 | 72 | 1 | 1 |  |  |  | 1.00 |
| 3 | 3_32 | 60 | 1 | 1 |  |  |  | 1.00 |
| 3 | 3_33 | 74 | 4 | 0.4 | 0.3 | 0.25 | 0.05 | 3.17 |
| 3 | 3_34 | 41 | 2 | 0.65 | 0.35 |  |  | 1.83 |
| 3 | 3_35 | 69 | 1 | 1 |  |  |  | 1.00 |
| 3 | 3_37 | 85 | 3 | 0.6 | 0.35 | 0.05 |  | 2.06 |
| 3 | 3_39 | 73 | 1 | 1 |  |  |  | 1.00 |
| 3 | 3_4 | 20 | 1 | 1 |  |  |  | 1.00 |
| 3 | 3_40 | 26 | 1 | 1 |  |  |  | 1.00 |
| 3 | 3_42 | 30 | 2 | 0.95 | 0.05 |  |  | 1.10 |
| 3 | 3_45 | 42 | 1 | 1 |  |  |  | 1.00 |
| 3 | 3_49 | 60 | 2 | 0.95 | 0.05 |  |  | 1.10 |
| 3 | 3_50 | 117 | 1 | 1 |  |  |  | 1.00 |
| 3 | 3_52 | 32 | 1 | 1 |  |  |  | 1.00 |
| 3 | 3_56 | 39 | 2 | 0.9 | 0.1 |  |  | 1.22 |
| 3 | 3_60 | 15 | 1 | 1 |  |  |  | 1.00 |
| 3 | 3_61 | 19 | 3 | 0.7 | 0.25 | 0.05 |  | 1.80 |
| 3 | 3_65 | 31 | 1 | 1 |  |  |  | 1.00 |
| 3 | 3_67 | 12 | 1 | 1 |  |  |  | 1.00 |
| 3 | 3_70 | 70 | 1 | 1 |  |  |  | 1.00 |
| 3 | 3_8 | 33 | 2 | 0.65 | 0.35 |  |  | 1.83 |
| 3 | 3_80 | 111 | 1 | 1 |  |  |  | 1.00 |
| 3 | 3_83 | 53 | 2 | 0.95 | 0.05 |  |  | 1.10 |
| 3 | 3_84 | 60 | 2 | 0.9 | 0.1 |  |  | 1.22 |
| 3 | 3_85 | 37 | 2 | 0.95 | 0.05 |  |  | 1.10 |
| 3 | 3_86 | 34 | 2 | 0.6 | 0.4 |  |  | 1.92 |
| 3 | 3_87 | 123 | 1 | 1 |  |  |  | 1.00 |
| 3 | 3_90 | 36 | 2 | 0.65 | 0.35 |  |  | 1.83 |
| 3 | 3_92 | 25 | 1 | 1 |  |  |  | 1.00 |
| 3 | 3_95 | 36 | 1 | 1 |  |  |  | 1.00 |
| 3 | 3_96 | 35 | 2 | 0.65 | 0.35 |  |  | 1.83 |
| 3 | 3_97 | 84 | 1 | 1 |  |  |  | 1.00 |
| 4 | 4_1 | 37 | 1 | 1 |  |  |  | 1.00 |
| 4 | 4_10 | 49 | 2 | 0.95 | 0.05 |  |  | 1.10 |
| 4 | 4_100 | 63 | 2 | 0.95 | 0.05 |  |  | 1.10 |
| 4 | 4_105 | 53 | 2 | 0.95 | 0.05 |  |  | 1.10 |
| 4 | 4_106 | 12 | 1 | 1 |  |  |  | 1.00 |
| 4 | 4_107 | 41 | 1 | 1 |  |  |  | 1.00 |
| 4 | 4_11 | 50 | 1 | 1 |  |  |  | 1.00 |
| 4 | 4_110 | 64 | 2 | 0.95 | 0.05 |  |  | 1.10 |
| 4 | 4_111 | 13 | 1 | 1 |  |  |  | 1.00 |
| 4 | 4_123 | 60 | 1 | 1 |  |  |  | 1.00 |
| 4 | 4_125 | 97 | 2 | 0.95 | 0.05 |  |  | 1.10 |
| 4 | 4_126 | 37 | 1 | 1 |  |  |  | 1.00 |
| 4 | 4_127 | 12 | 1 | 1 |  |  |  | 1.00 |
| 4 | 4_128 | 72 | 1 | 1 |  |  |  | 1.00 |
| 4 | 4_15 | 81 | 1 | 1 |  |  |  | 1.00 |
| 4 | 4_18 | 37 | 1 | 1 |  |  |  | 1.00 |
| 4 | 4_25 | 71 | 1 | 1 |  |  |  | 1.00 |
| 4 | 4_27 | 48 | 1 | 1 |  |  |  | 1.00 |
| 4 | 4_28 | 56 | 1 | 1 |  |  |  | 1.00 |
| 4 | 4_3 | 67 | 1 | 1 |  |  |  | 1.00 |
| 4 | 4_33 | 66 | 2 | 0.95 | 0.05 |  |  | 1.10 |
| 4 | 4_34 | 32 | 1 | 1 |  |  |  | 1.00 |
| 4 | 4_36 | 25 | 1 | 1 |  |  |  | 1.00 |
| 4 | 4_37 | 69 | 1 | 1 |  |  |  | 1.00 |
| 4 | 4_38 | 61 | 1 | 1 |  |  |  | 1.00 |
| 4 | 4_39 | 68 | 1 | 1 |  |  |  | 1.00 |
| 4 | 4_40 | 82 | 1 | 1 |  |  |  | 1.00 |
| 4 | 4_44 | 37 | 1 | 1 |  |  |  | 1.00 |
| 4 | 4_45 | 22 | 1 | 1 |  |  |  | 1.00 |
| 4 | 4_47 | 92 | 1 | 1 |  |  |  | 1.00 |
| 4 | 4_53 | 31 | 1 | 1 |  |  |  | 1.00 |
| 4 | 4_54 | 44 | 1 | 1 |  |  |  | 1.00 |
| 4 | 4_55 | 43 | 1 | 1 |  |  |  | 1.00 |
| 4 | 4_56 | 55 | 1 | 1 |  |  |  | 1.00 |
| 4 | 4_59 | 53 | 1 | 1 |  |  |  | 1.00 |
| 4 | 4_60 | 36 | 1 | 1 |  |  |  | 1.00 |
| 4 | 4_61 | 10 | 1 | 1 |  |  |  | 1.00 |
| 4 | 4_63 | 59 | 1 | 1 |  |  |  | 1.00 |
| 4 | 4_65 | 137 | 1 | 1 |  |  |  | 1.00 |
| 4 | 4_66 | 25 | 1 | 1 |  |  |  | 1.00 |
| 4 | 4_67 | 80 | 1 | 1 |  |  |  | 1.00 |
| 4 | 4_68 | 31 | 1 | 1 |  |  |  | 1.00 |
| 4 | 4_69 | 63 | 1 | 1 |  |  |  | 1.00 |
| 4 | 4_7 | 59 | 1 | 1 |  |  |  | 1.00 |
| 4 | 4_70 | 40 | 1 | 1 |  |  |  | 1.00 |
| 4 | 4_72 | 77 | 1 | 1 |  |  |  | 1.00 |
| 4 | 4_75 | 15 | 1 | 1 |  |  |  | 1.00 |
| 4 | 4_78 | 29 | 2 | 0.8 | 0.2 |  |  | 1.47 |
| 4 | 4_79 | 72 | 1 | 1 |  |  |  | 1.00 |
| 4 | 4_8 | 46 | 1 | 1 |  |  |  | 1.00 |
| 4 | 4_80 | 48 | 1 | 1 |  |  |  | 1.00 |
| 4 | 4_84 | 56 | 1 | 1 |  |  |  | 1.00 |
| 4 | 4_87 | 58 | 2 | 0.5 | 0.5 |  |  | 2.00 |
| 4 | 4_89 | 25 | 1 | 1 |  |  |  | 1.00 |
| 4 | 4_90 | 61 | 1 | 1 |  |  |  | 1.00 |
| 4 | 4_91 | 36 | 1 | 1 |  |  |  | 1.00 |
| 4 | 4_92 | 87 | 1 | 1 |  |  |  | 1.00 |
| 4 | 4_94 | 62 | 1 | 1 |  |  |  | 1.00 |
| 4 | 4_96 | 77 | 1 | 1 |  |  |  | 1.00 |
| 4 | 4_97 | 80 | 1 | 1 |  |  |  | 1.00 |
| 4 | 4_98 | 38 | 1 | 1 |  |  |  | 1.00 |
| 4 | 4_99 | 65 | 2 | 0.95 | 0.05 |  |  | 1.10 |

**Supplemental Table S2. The estimated polyandry for each family is reported when using only bi-allelic loci. All conventions arethe same as in Table S1.**

| Collection | Family | Number offspring | No Sires | Proportions | | | | Effective number sires |
| --- | --- | --- | --- | --- | --- | --- | --- | --- |
|  |  |  |  | w1 | w2 | w3 | w4 |  |
| 1 | 1_101 | 16 | 2 | 0.65 | 0.35 |  |  | 1.83 |
| 1 | 1_102 | 40 | 2 | 0.85 | 0.15 |  |  | 1.34 |
| 1 | 1_105 | 11 | 1 | 1 |  |  |  | 1 |
| 1 | 1_108 | 21 | 1 | 1 |  |  |  | 1 |
| 1 | 1_11 | 20 | 2 | 0.95 | 0.05 |  |  | 1.1 |
| 1 | 1_111 | 25 | 2 | 0.75 | 0.25 |  |  | 1.6 |
| 1 | 1_116 | 79 | 2 | 0.95 | 0.05 |  |  | 1.1 |
| 1 | 1_12 | 104 | 3 | 0.9 | 0.05 | 0.05 |  | 1.23 |
| 1 | 1_13 | 36 | 1 | 1 |  |  |  | 1 |
| 1 | 1_19 | 41 | 2 | 0.95 | 0.05 |  |  | 1.1 |
| 1 | 1_20 | 24 | 1 | 1 |  |  |  | 1 |
| 1 | 1_24 | 28 | 1 | 1 |  |  |  | 1 |
| 1 | 1_25 | 34 | 1 | 1 |  |  |  | 1 |
| 1 | 1_26 | 87 | 2 | 0.95 | 0.05 |  |  | 1.1 |
| 1 | 1_27 | 39 | 1 | 1 |  |  |  | 1 |
| 1 | 1_28 | 59 | 2 | 0.95 | 0.05 |  |  | 1.1 |
| 1 | 1_31 | 46 | 2 | 0.95 | 0.05 |  |  | 1.1 |
| 1 | 1_33 | 71 | 2 | 0.9 | 0.1 |  |  | 1.22 |
| 1 | 1_38 | 39 | 1 | 1 |  |  |  | 1 |
| 1 | 1_39 | 62 | 2 | 0.95 | 0.05 |  |  | 1.1 |
| 1 | 1_4 | 11 | 2 | 0.85 | 0.15 |  |  | 1.34 |
| 1 | 1_40 | 72 | 2 | 0.95 | 0.05 |  |  | 1.1 |
| 1 | 1_42 | 53 | 1 | 1 |  |  |  | 1 |
| 1 | 1_45 | 57 | 2 | 0.95 | 0.05 |  |  | 1.1 |
| 1 | 1_46 | 43 | 2 | 0.5 | 0.5 |  |  | 2 |
| 1 | 1_47 | 58 | 2 | 0.9 | 0.1 |  |  | 1.22 |
| 1 | 1_5 | 32 | 1 | 1 |  |  |  | 1 |
| 1 | 1_56 | 17 | 1 | 1 |  |  |  | 1 |
| 1 | 1_58 | 38 | 2 | 0.95 | 0.05 |  |  | 1.1 |
| 1 | 1_61 | 44 | 2 | 0.85 | 0.15 |  |  | 1.34 |
| 1 | 1_63 | 77 | 1 | 1 |  |  |  | 1 |
| 1 | 1_64 | 29 | 2 | 0.95 | 0.05 |  |  | 1.1 |
| 1 | 1_65 | 22 | 2 | 0.8 | 0.2 |  |  | 1.47 |
| 1 | 1_67 | 23 | 1 | 1 |  |  |  | 1 |
| 1 | 1_69 | 49 | 2 | 0.95 | 0.05 |  |  | 1.1 |
| 1 | 1_70 | 21 | 3 | 0.85 | 0.1 | 0.05 |  | 1.36 |
| 1 | 1_71 | 30 | 2 | 0.5 | 0.5 |  |  | 2 |
| 1 | 1_72 | 83 | 3 | 0.65 | 0.3 | 0.05 |  | 1.94 |
| 1 | 1_73 | 35 | 2 | 0.75 | 0.25 |  |  | 1.6 |
| 1 | 1_74 | 59 | 2 | 0.85 | 0.15 |  |  | 1.34 |
| 1 | 1_75 | 39 | 2 | 0.9 | 0.1 |  |  | 1.22 |
| 1 | 1_76 | 47 | 2 | 0.65 | 0.35 |  |  | 1.83 |
| 1 | 1_79 | 64 | 1 | 1 |  |  |  | 1 |
| 1 | 1_8 | 32 | 1 | 1 |  |  |  | 1 |
| 1 | 1_80 | 43 | 1 | 1 |  |  |  | 1 |
| 1 | 1_83 | 44 | 1 | 1 |  |  |  | 1 |
| 1 | 1_86 | 38 | 2 | 0.9 | 0.1 |  |  | 1.22 |
| 1 | 1_88 | 29 | 1 | 1 |  |  |  | 1 |
| 1 | 1_89 | 21 | 1 | 1 |  |  |  | 1 |
| 1 | 1_90 | 64 | 2 | 0.95 | 0.05 |  |  | 1.1 |
| 1 | 1_91 | 20 | 1 | 1 |  |  |  | 1 |
| 1 | 1_93 | 68 | 1 | 1 |  |  |  | 1 |
| 1 | 1_94 | 31 | 2 | 0.95 | 0.05 |  |  | 1.1 |
| 1 | 1_97 | 67 | 1 | 1 |  |  |  | 1 |
| 2 | 2_10 | 82 | 3 | 0.9 | 0.05 | 0.05 |  | 1.23 |
| 2 | 2_100 | 92 | 2 | 0.8 | 0.2 |  |  | 1.47 |
| 2 | 2_101 | 41 | 1 | 1 |  |  |  | 1 |
| 2 | 2_102 | 62 | 2 | 0.9 | 0.1 |  |  | 1.22 |
| 2 | 2_103 | 33 | 1 | 1 |  |  |  | 1 |
| 2 | 2_105 | 49 | 2 | 0.85 | 0.15 |  |  | 1.34 |
| 2 | 2_108 | 61 | 1 | 1 |  |  |  | 1 |
| 2 | 2_12 | 66 | 1 | 1 |  |  |  | 1 |
| 2 | 2_122 | 58 | 2 | 0.95 | 0.05 |  |  | 1.1 |
| 2 | 2_123 | 30 | 2 | 0.75 | 0.25 |  |  | 1.6 |
| 2 | 2_125 | 24 | 1 | 1 |  |  |  | 1 |
| 2 | 2_127 | 26 | 3 | 0.45 | 0.45 | 0.1 |  | 2.41 |
| 2 | 2_132 | 42 | 2 | 0.95 | 0.05 |  |  | 1.1 |
| 2 | 2_136 | 43 | 2 | 0.75 | 0.25 |  |  | 1.6 |
| 2 | 2_137 | 83 | 1 | 1 |  |  |  | 1 |
| 2 | 2_14 | 44 | 2 | 0.85 | 0.15 |  |  | 1.34 |
| 2 | 2_16 | 65 | 1 | 1 |  |  |  | 1 |
| 2 | 2_2 | 64 | 3 | 0.9 | 0.05 | 0.05 |  | 1.23 |
| 2 | 2_25 | 81 | 1 | 1 |  |  |  | 1 |
| 2 | 2_26 | 59 | 2 | 0.95 | 0.05 |  |  | 1.1 |
| 2 | 2_27 | 78 | 2 | 0.65 | 0.35 |  |  | 1.83 |
| 2 | 2_28 | 71 | 1 | 1 |  |  |  | 1 |
| 2 | 2_29 | 43 | 2 | 0.75 | 0.25 |  |  | 1.6 |
| 2 | 2_3 | 72 | 2 | 0.8 | 0.2 |  |  | 1.47 |
| 2 | 2_30 | 49 | 2 | 0.95 | 0.05 |  |  | 1.1 |
| 2 | 2_32 | 58 | 2 | 0.95 | 0.05 |  |  | 1.1 |
| 2 | 2_33 | 51 | 2 | 0.95 | 0.05 |  |  | 1.1 |
| 2 | 2_34 | 54 | 2 | 0.95 | 0.05 |  |  | 1.1 |
| 2 | 2_35 | 86 | 3 | 0.5 | 0.45 | 0.05 |  | 2.2 |
| 2 | 2_36 | 63 | 2 | 0.95 | 0.05 |  |  | 1.1 |
| 2 | 2_38 | 58 | 2 | 0.95 | 0.05 |  |  | 1.1 |
| 2 | 2_39 | 61 | 1 | 1 |  |  |  | 1 |
| 2 | 2_42 | 39 | 1 | 1 |  |  |  | 1 |
| 2 | 2_43 | 80 | 2 | 0.95 | 0.05 |  |  | 1.1 |
| 2 | 2_45 | 84 | 2 | 0.95 | 0.05 |  |  | 1.1 |
| 2 | 2_49 | 72 | 1 | 1 |  |  |  | 1 |
| 2 | 2_50 | 74 | 1 | 1 |  |  |  | 1 |
| 2 | 2_51 | 32 | 1 | 1 |  |  |  | 1 |
| 2 | 2_52 | 107 | 2 | 0.95 | 0.05 |  |  | 1.1 |
| 2 | 2_53 | 75 | 2 | 0.7 | 0.3 |  |  | 1.72 |
| 2 | 2_54 | 41 | 1 | 1 |  |  |  | 1 |
| 2 | 2_55 | 66 | 2 | 0.95 | 0.05 |  |  | 1.1 |
| 2 | 2_57 | 115 | 2 | 0.9 | 0.1 |  |  | 1.22 |
| 2 | 2_58 | 86 | 2 | 0.95 | 0.05 |  |  | 1.1 |
| 2 | 2_59 | 76 | 2 | 0.95 | 0.05 |  |  | 1.1 |
| 2 | 2_6 | 38 | 1 | 1 |  |  |  | 1 |
| 2 | 2_64 | 90 | 1 | 1 |  |  |  | 1 |
| 2 | 2_66 | 90 | 1 | 1 |  |  |  | 1 |
| 2 | 2_68 | 59 | 2 | 0.9 | 0.1 |  |  | 1.22 |
| 2 | 2_69 | 52 | 2 | 0.95 | 0.05 |  |  | 1.1 |
| 2 | 2_7 | 21 | 2 | 0.7 | 0.3 |  |  | 1.72 |
| 2 | 2_70 | 31 | 2 | 0.95 | 0.05 |  |  | 1.1 |
| 2 | 2_71 | 78 | 2 | 0.95 | 0.05 |  |  | 1.1 |
| 2 | 2_72 | 34 | 1 | 1 |  |  |  | 1 |
| 2 | 2_73 | 91 | 1 | 1 |  |  |  | 1 |
| 2 | 2_75 | 74 | 2 | 0.95 | 0.05 |  |  | 1.1 |
| 2 | 2_76 | 93 | 2 | 0.8 | 0.2 |  |  | 1.47 |
| 2 | 2_78 | 30 | 2 | 0.6 | 0.4 |  |  | 1.92 |
| 2 | 2_79 | 103 | 3 | 0.9 | 0.05 | 0.05 |  | 1.23 |
| 2 | 2_80 | 91 | 1 | 1 |  |  |  | 1 |
| 2 | 2_83 | 27 | 2 | 0.8 | 0.2 |  |  | 1.47 |
| 2 | 2_84 | 41 | 1 | 1 |  |  |  | 1 |
| 2 | 2_87 | 48 | 2 | 0.7 | 0.3 |  |  | 1.72 |
| 2 | 2_88 | 35 | 2 | 0.85 | 0.15 |  |  | 1.34 |
| 2 | 2_92 | 108 | 1 | 1 |  |  |  | 1 |
| 2 | 2_93 | 37 | 1 | 1 |  |  |  | 1 |
| 2 | 2_94 | 38 | 2 | 0.85 | 0.15 |  |  | 1.34 |
| 2 | 2_95 | 132 | 1 | 1 |  |  |  | 1 |
| 2 | 2_96 | 110 | 2 | 0.95 | 0.05 |  |  | 1.1 |
| 2 | 2_98 | 41 | 2 | 0.95 | 0.05 |  |  | 1.1 |
| 3 | 3_1 | 65 | 1 | 1 |  |  |  | 1 |
| 3 | 3_102 | 65 | 2 | 0.8 | 0.2 |  |  | 1.47 |
| 3 | 3_107 | 23 | 1 | 1 |  |  |  | 1 |
| 3 | 3_108 | 73 | 2 | 0.95 | 0.05 |  |  | 1.1 |
| 3 | 3_111 | 66 | 1 | 1 |  |  |  | 1 |
| 3 | 3_113 | 21 | 2 | 0.85 | 0.15 |  |  | 1.34 |
| 3 | 3_115 | 36 | 2 | 0.9 | 0.1 |  |  | 1.22 |
| 3 | 3_116 | 68 | 1 | 1 |  |  |  | 1 |
| 3 | 3_117 | 37 | 1 | 1 |  |  |  | 1 |
| 3 | 3_118 | 71 | 1 | 1 |  |  |  | 1 |
| 3 | 3_119 | 48 | 1 | 1 |  |  |  | 1 |
| 3 | 3_121 | 57 | 2 | 0.75 | 0.25 |  |  | 1.6 |
| 3 | 3_122 | 74 | 1 | 1 |  |  |  | 1 |
| 3 | 3_126 | 22 | 1 | 1 |  |  |  | 1 |
| 3 | 3_128 | 41 | 1 | 1 |  |  |  | 1 |
| 3 | 3_129 | 28 | 2 | 0.8 | 0.2 |  |  | 1.47 |
| 3 | 3_130 | 26 | 1 | 1 |  |  |  | 1 |
| 3 | 3_131 | 82 | 1 | 1 |  |  |  | 1 |
| 3 | 3_132 | 30 | 1 | 1 |  |  |  | 1 |
| 3 | 3_133 | 34 | 1 | 1 |  |  |  | 1 |
| 3 | 3_134 | 14 | 1 | 1 |  |  |  | 1 |
| 3 | 3_135 | 55 | 1 | 1 |  |  |  | 1 |
| 3 | 3_136 | 87 | 1 | 1 |  |  |  | 1 |
| 3 | 3_140 | 30 | 2 | 0.95 | 0.05 |  |  | 1.1 |
| 3 | 3_142 | 58 | 2 | 0.95 | 0.05 |  |  | 1.1 |
| 3 | 3_144 | 51 | 1 | 1 |  |  |  | 1 |
| 3 | 3_147 | 20 | 1 | 1 |  |  |  | 1 |
| 3 | 3_148 | 97 | 3 | 0.55 | 0.35 | 0.1 |  | 2.3 |
| 3 | 3_149 | 75 | 1 | 1 |  |  |  | 1 |
| 3 | 3_152 | 29 | 2 | 0.8 | 0.2 |  |  | 1.47 |
| 3 | 3_154 | 42 | 1 | 1 |  |  |  | 1 |
| 3 | 3_155 | 42 | 1 | 1 |  |  |  | 1 |
| 3 | 3_156 | 57 | 2 | 0.95 | 0.05 |  |  | 1.1 |
| 3 | 3_157 | 19 | 2 | 0.95 | 0.05 |  |  | 1.1 |
| 3 | 3_16 | 100 | 1 | 1 |  |  |  | 1 |
| 3 | 3_161 | 41 | 1 | 1 |  |  |  | 1 |
| 3 | 3_163 | 75 | 2 | 0.7 | 0.3 |  |  | 1.72 |
| 3 | 3_165 | 41 | 1 | 1 |  |  |  | 1 |
| 3 | 3_166 | 36 | 2 | 0.95 | 0.05 |  |  | 1.1 |
| 3 | 3_167 | 81 | 1 | 1 |  |  |  | 1 |
| 3 | 3_168 | 67 | 1 | 1 |  |  |  | 1 |
| 3 | 3_17 | 68 | 2 | 0.95 | 0.05 |  |  | 1.1 |
| 3 | 3_171 | 80 | 1 | 1 |  |  |  | 1 |
| 3 | 3_172 | 104 | 2 | 0.95 | 0.05 |  |  | 1.1 |
| 3 | 3_173 | 8 | 1 | 1 |  |  |  | 1 |
| 3 | 3_174 | 35 | 1 | 1 |  |  |  | 1 |
| 3 | 3_175 | 13 | 1 | 1 |  |  |  | 1 |
| 3 | 3_18 | 43 | 2 | 0.8 | 0.2 |  |  | 1.47 |
| 3 | 3_181 | 28 | 1 | 1 |  |  |  | 1 |
| 3 | 3_182 | 47 | 1 | 1 |  |  |  | 1 |
| 3 | 3_184 | 44 | 1 | 1 |  |  |  | 1 |
| 3 | 3_20 | 73 | 1 | 1 |  |  |  | 1 |
| 3 | 3_21 | 54 | 1 | 1 |  |  |  | 1 |
| 3 | 3_24 | 55 | 1 | 1 |  |  |  | 1 |
| 3 | 3_25 | 38 | 1 | 1 |  |  |  | 1 |
| 3 | 3_26 | 30 | 2 | 0.85 | 0.15 |  |  | 1.34 |
| 3 | 3_3 | 38 | 1 | 1 |  |  |  | 1 |
| 3 | 3_31 | 72 | 1 | 1 |  |  |  | 1 |
| 3 | 3_32 | 60 | 1 | 1 |  |  |  | 1 |
| 3 | 3_33 | 74 | 4 | 0.4 | 0.35 | 0.2 | 0.05 | 3.08 |
| 3 | 3_34 | 41 | 2 | 0.65 | 0.35 |  |  | 1.83 |
| 3 | 3_35 | 69 | 1 | 1 |  |  |  | 1 |
| 3 | 3_37 | 85 | 3 | 0.6 | 0.35 | 0.05 |  | 2.06 |
| 3 | 3_39 | 73 | 1 | 1 |  |  |  | 1 |
| 3 | 3_4 | 20 | 1 | 1 |  |  |  | 1 |
| 3 | 3_40 | 26 | 1 | 1 |  |  |  | 1 |
| 3 | 3_42 | 30 | 2 | 0.95 | 0.05 |  |  | 1.1 |
| 3 | 3_45 | 42 | 1 | 1 |  |  |  | 1 |
| 3 | 3_49 | 60 | 2 | 0.95 | 0.05 |  |  | 1.1 |
| 3 | 3_50 | 117 | 1 | 1 |  |  |  | 1 |
| 3 | 3_52 | 32 | 1 | 1 |  |  |  | 1 |
| 3 | 3_56 | 39 | 2 | 0.9 | 0.1 |  |  | 1.22 |
| 3 | 3_60 | 15 | 1 | 1 |  |  |  | 1 |
| 3 | 3_61 | 19 | 3 | 0.75 | 0.2 | 0.05 |  | 1.65 |
| 3 | 3_65 | 31 | 1 | 1 |  |  |  | 1 |
| 3 | 3_67 | 12 | 1 | 1 |  |  |  | 1 |
| 3 | 3_70 | 70 | 1 | 1 |  |  |  | 1 |
| 3 | 3_8 | 33 | 2 | 0.65 | 0.35 |  |  | 1.83 |
| 3 | 3_80 | 111 | 1 | 1 |  |  |  | 1 |
| 3 | 3_83 | 53 | 2 | 0.95 | 0.05 |  |  | 1.1 |
| 3 | 3_84 | 60 | 2 | 0.9 | 0.1 |  |  | 1.22 |
| 3 | 3_85 | 37 | 2 | 0.95 | 0.05 |  |  | 1.1 |
| 3 | 3_86 | 34 | 2 | 0.65 | 0.35 |  |  | 1.83 |
| 3 | 3_87 | 123 | 1 | 1 |  |  |  | 1 |
| 3 | 3_90 | 36 | 2 | 0.65 | 0.35 |  |  | 1.83 |
| 3 | 3_92 | 25 | 1 | 1 |  |  |  | 1 |
| 3 | 3_95 | 36 | 1 | 1 |  |  |  | 1 |
| 3 | 3_96 | 35 | 2 | 0.6 | 0.4 |  |  | 1.92 |
| 3 | 3_97 | 84 | 1 | 1 |  |  |  | 1 |
| 4 | 4_1 | 37 | 1 | 1 |  |  |  | 1 |
| 4 | 4_10 | 49 | 2 | 0.95 | 0.05 |  |  | 1.1 |
| 4 | 4_100 | 63 | 2 | 0.95 | 0.05 |  |  | 1.1 |
| 4 | 4_105 | 53 | 2 | 0.95 | 0.05 |  |  | 1.1 |
| 4 | 4_106 | 12 | 1 | 1 |  |  |  | 1 |
| 4 | 4_107 | 41 | 1 | 1 |  |  |  | 1 |
| 4 | 4_11 | 50 | 1 | 1 |  |  |  | 1 |
| 4 | 4_110 | 64 | 2 | 0.95 | 0.05 |  |  | 1.1 |
| 4 | 4_111 | 13 | 1 | 1 |  |  |  | 1 |
| 4 | 4_123 | 60 | 2 | 0.95 | 0.05 |  |  | 1.1 |
| 4 | 4_125 | 97 | 2 | 0.95 | 0.05 |  |  | 1.1 |
| 4 | 4_126 | 37 | 1 | 1 |  |  |  | 1 |
| 4 | 4_127 | 12 | 1 | 1 |  |  |  | 1 |
| 4 | 4_128 | 72 | 1 | 1 |  |  |  | 1 |
| 4 | 4_15 | 81 | 1 | 1 |  |  |  | 1 |
| 4 | 4_18 | 37 | 1 | 1 |  |  |  | 1 |
| 4 | 4_25 | 71 | 1 | 1 |  |  |  | 1 |
| 4 | 4_27 | 48 | 1 | 1 |  |  |  | 1 |
| 4 | 4_28 | 56 | 1 | 1 |  |  |  | 1 |
| 4 | 4_3 | 67 | 1 | 1 |  |  |  | 1 |
| 4 | 4_33 | 66 | 2 | 0.95 | 0.05 |  |  | 1.1 |
| 4 | 4_34 | 32 | 1 | 1 |  |  |  | 1 |
| 4 | 4_36 | 25 | 1 | 1 |  |  |  | 1 |
| 4 | 4_37 | 69 | 1 | 1 |  |  |  | 1 |
| 4 | 4_38 | 61 | 1 | 1 |  |  |  | 1 |
| 4 | 4_39 | 68 | 1 | 1 |  |  |  | 1 |
| 4 | 4_40 | 82 | 1 | 1 |  |  |  | 1 |
| 4 | 4_44 | 37 | 1 | 1 |  |  |  | 1 |
| 4 | 4_45 | 22 | 1 | 1 |  |  |  | 1 |
| 4 | 4_47 | 92 | 1 | 1 |  |  |  | 1 |
| 4 | 4_53 | 31 | 1 | 1 |  |  |  | 1 |
| 4 | 4_54 | 44 | 1 | 1 |  |  |  | 1 |
| 4 | 4_55 | 43 | 1 | 1 |  |  |  | 1 |
| 4 | 4_56 | 55 | 1 | 1 |  |  |  | 1 |
| 4 | 4_59 | 53 | 1 | 1 |  |  |  | 1 |
| 4 | 4_60 | 36 | 1 | 1 |  |  |  | 1 |
| 4 | 4_61 | 10 | 1 | 1 |  |  |  | 1 |
| 4 | 4_63 | 59 | 2 | 0.95 | 0.05 |  |  | 1.1 |
| 4 | 4_65 | 137 | 1 | 1 |  |  |  | 1 |
| 4 | 4_66 | 25 | 1 | 1 |  |  |  | 1 |
| 4 | 4_67 | 80 | 1 | 1 |  |  |  | 1 |
| 4 | 4_68 | 31 | 1 | 1 |  |  |  | 1 |
| 4 | 4_69 | 63 | 1 | 1 |  |  |  | 1 |
| 4 | 4_7 | 59 | 1 | 1 |  |  |  | 1 |
| 4 | 4_70 | 40 | 1 | 1 |  |  |  | 1 |
| 4 | 4_72 | 77 | 1 | 1 |  |  |  | 1 |
| 4 | 4_75 | 15 | 1 | 1 |  |  |  | 1 |
| 4 | 4_78 | 29 | 2 | 0.8 | 0.2 |  |  | 1.47 |
| 4 | 4_79 | 72 | 1 | 1 |  |  |  | 1 |
| 4 | 4_8 | 46 | 1 | 1 |  |  |  | 1 |
| 4 | 4_80 | 48 | 1 | 1 |  |  |  | 1 |
| 4 | 4_84 | 56 | 1 | 1 |  |  |  | 1 |
| 4 | 4_87 | 58 | 2 | 0.5 | 0.5 |  |  | 2 |
| 4 | 4_89 | 25 | 1 | 1 |  |  |  | 1 |
| 4 | 4_90 | 61 | 1 | 1 |  |  |  | 1 |
| 4 | 4_91 | 36 | 1 | 1 |  |  |  | 1 |
| 4 | 4_92 | 87 | 1 | 1 |  |  |  | 1 |
| 4 | 4_94 | 62 | 1 | 1 |  |  |  | 1 |
| 4 | 4_96 | 77 | 1 | 1 |  |  |  | 1 |
| 4 | 4_97 | 80 | 1 | 1 |  |  |  | 1 |
| 4 | 4_98 | 38 | 1 | 1 |  |  |  | 1 |
| 4 | 4_99 | 65 | 2 | 0.95 | 0.05 |  |  | 1.1 |

**Supplemental Table S3. The estimated polyandry for each family is reported when using only 4-allele loci. All conventions are the same as in Table S1.**

| Collection | Family | Number offspring | No Sires | Proportions | | | | Effective number sires |
| --- | --- | --- | --- | --- | --- | --- | --- | --- |
|  |  |  |  | w1 | w2 | w3 | w4 |  |
| 1 | 1_101 | 16 | 2 | 0.65 | 0.35 |  |  | 1.83 |
| 1 | 1_102 | 40 | 2 | 0.85 | 0.15 |  |  | 1.34 |
| 1 | 1_105 | 11 | 1 | 1 |  |  |  | 1 |
| 1 | 1_108 | 21 | 1 | 1 |  |  |  | 1 |
| 1 | 1_11 | 20 | 2 | 0.9 | 0.1 |  |  | 1.22 |
| 1 | 1_111 | 25 | 2 | 0.7 | 0.3 |  |  | 1.72 |
| 1 | 1_116 | 79 | 2 | 0.9 | 0.1 |  |  | 1.22 |
| 1 | 1_12 | 104 | 2 | 0.95 | 0.05 |  |  | 1.1 |
| 1 | 1_13 | 36 | 1 | 1 |  |  |  | 1 |
| 1 | 1_19 | 41 | 2 | 0.95 | 0.05 |  |  | 1.1 |
| 1 | 1_20 | 24 | 1 | 1 |  |  |  | 1 |
| 1 | 1_24 | 28 | 1 | 1 |  |  |  | 1 |
| 1 | 1_25 | 34 | 1 | 1 |  |  |  | 1 |
| 1 | 1_26 | 87 | 2 | 0.95 | 0.05 |  |  | 1.1 |
| 1 | 1_27 | 39 | 1 | 1 |  |  |  | 1 |
| 1 | 1_28 | 59 | 2 | 0.95 | 0.05 |  |  | 1.1 |
| 1 | 1_31 | 46 | 1 | 1 |  |  |  | 1 |
| 1 | 1_33 | 71 | 2 | 0.9 | 0.1 |  |  | 1.22 |
| 1 | 1_38 | 39 | 1 | 1 |  |  |  | 1 |
| 1 | 1_39 | 62 | 2 | 0.95 | 0.05 |  |  | 1.1 |
| 1 | 1_4 | 11 | 2 | 0.8 | 0.2 |  |  | 1.47 |
| 1 | 1_40 | 72 | 1 | 1 |  |  |  | 1 |
| 1 | 1_42 | 53 | 1 | 1 |  |  |  | 1 |
| 1 | 1_45 | 57 | 1 | 1 |  |  |  | 1 |
| 1 | 1_46 | 43 | 2 | 0.6 | 0.4 |  |  | 1.92 |
| 1 | 1_47 | 58 | 2 | 0.9 | 0.1 |  |  | 1.22 |
| 1 | 1_5 | 32 | 1 | 1 |  |  |  | 1 |
| 1 | 1_56 | 17 | 1 | 1 |  |  |  | 1 |
| 1 | 1_58 | 38 | 1 | 1 |  |  |  | 1 |
| 1 | 1_61 | 44 | 2 | 0.85 | 0.15 |  |  | 1.34 |
| 1 | 1_63 | 77 | 1 | 1 |  |  |  | 1 |
| 1 | 1_64 | 29 | 2 | 0.95 | 0.05 |  |  | 1.1 |
| 1 | 1_65 | 22 | 2 | 0.8 | 0.2 |  |  | 1.47 |
| 1 | 1_67 | 23 | 1 | 1 |  |  |  | 1 |
| 1 | 1_69 | 49 | 2 | 0.95 | 0.05 |  |  | 1.1 |
| 1 | 1_70 | 21 | 2 | 0.85 | 0.15 |  |  | 1.34 |
| 1 | 1_71 | 30 | 2 | 0.6 | 0.4 |  |  | 1.92 |
| 1 | 1_72 | 83 | 3 | 0.6 | 0.3 | 0.1 |  | 2.17 |
| 1 | 1_73 | 35 | 2 | 0.7 | 0.3 |  |  | 1.72 |
| 1 | 1_74 | 59 | 2 | 0.9 | 0.1 |  |  | 1.22 |
| 1 | 1_75 | 39 | 2 | 0.9 | 0.1 |  |  | 1.22 |
| 1 | 1_76 | 47 | 2 | 0.65 | 0.35 |  |  | 1.83 |
| 1 | 1_79 | 64 | 1 | 1 |  |  |  | 1 |
| 1 | 1_8 | 32 | 1 | 1 |  |  |  | 1 |
| 1 | 1_80 | 43 | 1 | 1 |  |  |  | 1 |
| 1 | 1_83 | 44 | 1 | 1 |  |  |  | 1 |
| 1 | 1_86 | 38 | 2 | 0.9 | 0.1 |  |  | 1.22 |
| 1 | 1_88 | 29 | 1 | 1 |  |  |  | 1 |
| 1 | 1_89 | 21 | 1 | 1 |  |  |  | 1 |
| 1 | 1_90 | 64 | 2 | 0.95 | 0.05 |  |  | 1.1 |
| 1 | 1_91 | 20 | 1 | 1 |  |  |  | 1 |
| 1 | 1_93 | 68 | 1 | 1 |  |  |  | 1 |
| 1 | 1_94 | 31 | 2 | 0.9 | 0.1 |  |  | 1.22 |
| 1 | 1_97 | 67 | 1 | 1 |  |  |  | 1 |
| 2 | 2_10 | 82 | 2 | 0.95 | 0.05 |  |  | 1.1 |
| 2 | 2_100 | 92 | 2 | 0.8 | 0.2 |  |  | 1.47 |
| 2 | 2_101 | 41 | 1 | 1 |  |  |  | 1 |
| 2 | 2_102 | 62 | 2 | 0.9 | 0.1 |  |  | 1.22 |
| 2 | 2_103 | 33 | 1 | 1 |  |  |  | 1 |
| 2 | 2_105 | 49 | 2 | 0.85 | 0.15 |  |  | 1.34 |
| 2 | 2_108 | 61 | 1 | 1 |  |  |  | 1 |
| 2 | 2_12 | 66 | 1 | 1 |  |  |  | 1 |
| 2 | 2_122 | 58 | 2 | 0.95 | 0.05 |  |  | 1.1 |
| 2 | 2_123 | 30 | 2 | 0.8 | 0.2 |  |  | 1.47 |
| 2 | 2_125 | 24 | 1 | 1 |  |  |  | 1 |
| 2 | 2_127 | 26 | 2 | 0.7 | 0.3 |  |  | 1.72 |
| 2 | 2_132 | 42 | 2 | 0.95 | 0.05 |  |  | 1.1 |
| 2 | 2_136 | 43 | 2 | 0.75 | 0.25 |  |  | 1.6 |
| 2 | 2_137 | 83 | 1 | 1 |  |  |  | 1 |
| 2 | 2_14 | 44 | 2 | 0.9 | 0.1 |  |  | 1.22 |
| 2 | 2_16 | 65 | 1 | 1 |  |  |  | 1 |
| 2 | 2_2 | 64 | 3 | 0.85 | 0.1 | 0.05 |  | 1.36 |
| 2 | 2_25 | 81 | 1 | 1 |  |  |  | 1 |
| 2 | 2_26 | 59 | 1 | 1 |  |  |  | 1 |
| 2 | 2_27 | 78 | 2 | 0.65 | 0.35 |  |  | 1.83 |
| 2 | 2_28 | 71 | 1 | 1 |  |  |  | 1 |
| 2 | 2_29 | 43 | 1 | 1 |  |  |  | 1 |
| 2 | 2_3 | 72 | 2 | 0.85 | 0.15 |  |  | 1.34 |
| 2 | 2_30 | 49 | 1 | 1 |  |  |  | 1 |
| 2 | 2_32 | 58 | 1 | 1 |  |  |  | 1 |
| 2 | 2_33 | 51 | 1 | 1 |  |  |  | 1 |
| 2 | 2_34 | 54 | 2 | 0.95 | 0.05 |  |  | 1.1 |
| 2 | 2_35 | 86 | 3 | 0.55 | 0.4 | 0.05 |  | 2.15 |
| 2 | 2_36 | 63 | 2 | 0.95 | 0.05 |  |  | 1.1 |
| 2 | 2_38 | 58 | 2 | 0.95 | 0.05 |  |  | 1.1 |
| 2 | 2_39 | 61 | 1 | 1 |  |  |  | 1 |
| 2 | 2_42 | 39 | 1 | 1 |  |  |  | 1 |
| 2 | 2_43 | 80 | 2 | 0.95 | 0.05 |  |  | 1.1 |
| 2 | 2_45 | 84 | 2 | 0.95 | 0.05 |  |  | 1.1 |
| 2 | 2_49 | 72 | 1 | 1 |  |  |  | 1 |
| 2 | 2_50 | 74 | 1 | 1 |  |  |  | 1 |
| 2 | 2_51 | 32 | 1 | 1 |  |  |  | 1 |
| 2 | 2_52 | 107 | 2 | 0.95 | 0.05 |  |  | 1.1 |
| 2 | 2_53 | 75 | 2 | 0.7 | 0.3 |  |  | 1.72 |
| 2 | 2_54 | 41 | 1 | 1 |  |  |  | 1 |
| 2 | 2_55 | 66 | 2 | 0.95 | 0.05 |  |  | 1.1 |
| 2 | 2_57 | 115 | 2 | 0.9 | 0.1 |  |  | 1.22 |
| 2 | 2_58 | 86 | 2 | 0.95 | 0.05 |  |  | 1.1 |
| 2 | 2_59 | 76 | 2 | 0.95 | 0.05 |  |  | 1.1 |
| 2 | 2_6 | 38 | 1 | 1 |  |  |  | 1 |
| 2 | 2_64 | 90 | 1 | 1 |  |  |  | 1 |
| 2 | 2_66 | 90 | 1 | 1 |  |  |  | 1 |
| 2 | 2_68 | 59 | 2 | 0.9 | 0.1 |  |  | 1.22 |
| 2 | 2_69 | 52 | 2 | 0.95 | 0.05 |  |  | 1.1 |
| 2 | 2_7 | 21 | 2 | 0.75 | 0.25 |  |  | 1.6 |
| 2 | 2_70 | 31 | 2 | 0.95 | 0.05 |  |  | 1.1 |
| 2 | 2_71 | 78 | 2 | 0.95 | 0.05 |  |  | 1.1 |
| 2 | 2_72 | 34 | 1 | 1 |  |  |  | 1 |
| 2 | 2_73 | 91 | 1 | 1 |  |  |  | 1 |
| 2 | 2_75 | 74 | 2 | 0.95 | 0.05 |  |  | 1.1 |
| 2 | 2_76 | 93 | 2 | 0.85 | 0.15 |  |  | 1.34 |
| 2 | 2_78 | 30 | 2 | 0.65 | 0.35 |  |  | 1.83 |
| 2 | 2_79 | 103 | 2 | 0.9 | 0.1 |  |  | 1.22 |
| 2 | 2_80 | 91 | 1 | 1 |  |  |  | 1 |
| 2 | 2_83 | 27 | 2 | 0.8 | 0.2 |  |  | 1.47 |
| 2 | 2_84 | 41 | 1 | 1 |  |  |  | 1 |
| 2 | 2_87 | 48 | 2 | 0.7 | 0.3 |  |  | 1.72 |
| 2 | 2_88 | 35 | 2 | 0.85 | 0.15 |  |  | 1.34 |
| 2 | 2_92 | 108 | 1 | 1 |  |  |  | 1 |
| 2 | 2_93 | 37 | 1 | 1 |  |  |  | 1 |
| 2 | 2_94 | 38 | 2 | 0.85 | 0.15 |  |  | 1.34 |
| 2 | 2_95 | 132 | 1 | 1 |  |  |  | 1 |
| 2 | 2_96 | 110 | 2 | 0.95 | 0.05 |  |  | 1.1 |
| 2 | 2_98 | 41 | 2 | 0.95 | 0.05 |  |  | 1.1 |
| 3 | 3_1 | 65 | 1 | 1 |  |  |  | 1 |
| 3 | 3_102 | 65 | 1 | 1 |  |  |  | 1 |
| 3 | 3_107 | 23 | 1 | 1 |  |  |  | 1 |
| 3 | 3_108 | 73 | 2 | 0.95 | 0.05 |  |  | 1.1 |
| 3 | 3_111 | 66 | 1 | 1 |  |  |  | 1 |
| 3 | 3_113 | 21 | 2 | 0.9 | 0.1 |  |  | 1.22 |
| 3 | 3_115 | 36 | 2 | 0.9 | 0.1 |  |  | 1.22 |
| 3 | 3_116 | 68 | 1 | 1 |  |  |  | 1 |
| 3 | 3_117 | 37 | 1 | 1 |  |  |  | 1 |
| 3 | 3_118 | 71 | 1 | 1 |  |  |  | 1 |
| 3 | 3_119 | 48 | 1 | 1 |  |  |  | 1 |
| 3 | 3_121 | 57 | 2 | 0.7 | 0.3 |  |  | 1.72 |
| 3 | 3_122 | 74 | 1 | 1 |  |  |  | 1 |
| 3 | 3_126 | 22 | 1 | 1 |  |  |  | 1 |
| 3 | 3_128 | 41 | 1 | 1 |  |  |  | 1 |
| 3 | 3_129 | 28 | 2 | 0.8 | 0.2 |  |  | 1.47 |
| 3 | 3_130 | 26 | 1 | 1 |  |  |  | 1 |
| 3 | 3_131 | 82 | 1 | 1 |  |  |  | 1 |
| 3 | 3_132 | 30 | 1 | 1 |  |  |  | 1 |
| 3 | 3_133 | 34 | 1 | 1 |  |  |  | 1 |
| 3 | 3_134 | 14 | 1 | 1 |  |  |  | 1 |
| 3 | 3_135 | 55 | 1 | 1 |  |  |  | 1 |
| 3 | 3_136 | 87 | 1 | 1 |  |  |  | 1 |
| 3 | 3_140 | 30 | 2 | 0.95 | 0.05 |  |  | 1.1 |
| 3 | 3_142 | 58 | 2 | 0.95 | 0.05 |  |  | 1.1 |
| 3 | 3_144 | 51 | 1 | 1 |  |  |  | 1 |
| 3 | 3_147 | 20 | 1 | 1 |  |  |  | 1 |
| 3 | 3_148 | 97 | 3 | 0.5 | 0.4 | 0.1 |  | 2.38 |
| 3 | 3_149 | 75 | 1 | 1 |  |  |  | 1 |
| 3 | 3_152 | 29 | 2 | 0.75 | 0.25 |  |  | 1.6 |
| 3 | 3_154 | 42 | 1 | 1 |  |  |  | 1 |
| 3 | 3_155 | 42 | 1 | 1 |  |  |  | 1 |
| 3 | 3_156 | 57 | 1 | 1 |  |  |  | 1 |
| 3 | 3_157 | 19 | 2 | 0.9 | 0.1 |  |  | 1.22 |
| 3 | 3_16 | 100 | 1 | 1 |  |  |  | 1 |
| 3 | 3_161 | 41 | 1 | 1 |  |  |  | 1 |
| 3 | 3_163 | 75 | 1 | 1 |  |  |  | 1 |
| 3 | 3_165 | 41 | 1 | 1 |  |  |  | 1 |
| 3 | 3_166 | 36 | 2 | 0.95 | 0.05 |  |  | 1.1 |
| 3 | 3_167 | 81 | 1 | 1 |  |  |  | 1 |
| 3 | 3_168 | 67 | 1 | 1 |  |  |  | 1 |
| 3 | 3_17 | 68 | 1 | 1 |  |  |  | 1 |
| 3 | 3_171 | 80 | 1 | 1 |  |  |  | 1 |
| 3 | 3_172 | 104 | 2 | 0.95 | 0.05 |  |  | 1.1 |
| 3 | 3_173 | 8 | 1 | 1 |  |  |  | 1 |
| 3 | 3_174 | 35 | 1 | 1 |  |  |  | 1 |
| 3 | 3_175 | 13 | 1 | 1 |  |  |  | 1 |
| 3 | 3_18 | 43 | 2 | 0.8 | 0.2 |  |  | 1.47 |
| 3 | 3_181 | 28 | 1 | 1 |  |  |  | 1 |
| 3 | 3_182 | 47 | 1 | 1 |  |  |  | 1 |
| 3 | 3_184 | 44 | 1 | 1 |  |  |  | 1 |
| 3 | 3_20 | 73 | 1 | 1 |  |  |  | 1 |
| 3 | 3_21 | 54 | 1 | 1 |  |  |  | 1 |
| 3 | 3_24 | 55 | 1 | 1 |  |  |  | 1 |
| 3 | 3_25 | 38 | 1 | 1 |  |  |  | 1 |
| 3 | 3_26 | 30 | 2 | 0.8 | 0.2 |  |  | 1.47 |
| 3 | 3_3 | 38 | 1 | 1 |  |  |  | 1 |
| 3 | 3_31 | 72 | 1 | 1 |  |  |  | 1 |
| 3 | 3_32 | 60 | 1 | 1 |  |  |  | 1 |
| 3 | 3_33 | 74 | 4 | 0.4 | 0.3 | 0.25 | 0.05 | 3.17 |
| 3 | 3_34 | 41 | 2 | 0.7 | 0.3 |  |  | 1.72 |
| 3 | 3_35 | 69 | 1 | 1 |  |  |  | 1 |
| 3 | 3_37 | 85 | 3 | 0.6 | 0.35 | 0.05 |  | 2.06 |
| 3 | 3_39 | 73 | 1 | 1 |  |  |  | 1 |
| 3 | 3_4 | 20 | 1 | 1 |  |  |  | 1 |
| 3 | 3_40 | 26 | 1 | 1 |  |  |  | 1 |
| 3 | 3_42 | 30 | 2 | 0.95 | 0.05 |  |  | 1.1 |
| 3 | 3_45 | 42 | 1 | 1 |  |  |  | 1 |
| 3 | 3_49 | 60 | 2 | 0.95 | 0.05 |  |  | 1.1 |
| 3 | 3_50 | 117 | 1 | 1 |  |  |  | 1 |
| 3 | 3_52 | 32 | 1 | 1 |  |  |  | 1 |
| 3 | 3_56 | 39 | 2 | 0.9 | 0.1 |  |  | 1.22 |
| 3 | 3_60 | 15 | 1 | 1 |  |  |  | 1 |
| 3 | 3_61 | 19 | 3 | 0.6 | 0.35 | 0.05 |  | 2.06 |
| 3 | 3_65 | 31 | 1 | 1 |  |  |  | 1 |
| 3 | 3_67 | 12 | 1 | 1 |  |  |  | 1 |
| 3 | 3_70 | 70 | 1 | 1 |  |  |  | 1 |
| 3 | 3_8 | 33 | 2 | 0.7 | 0.3 |  |  | 1.72 |
| 3 | 3_80 | 111 | 1 | 1 |  |  |  | 1 |
| 3 | 3_83 | 53 | 2 | 0.95 | 0.05 |  |  | 1.1 |
| 3 | 3_84 | 60 | 2 | 0.9 | 0.1 |  |  | 1.22 |
| 3 | 3_85 | 37 | 2 | 0.95 | 0.05 |  |  | 1.1 |
| 3 | 3_86 | 34 | 2 | 0.6 | 0.4 |  |  | 1.92 |
| 3 | 3_87 | 123 | 1 | 1 |  |  |  | 1 |
| 3 | 3_90 | 36 | 2 | 0.65 | 0.35 |  |  | 1.83 |
| 3 | 3_92 | 25 | 1 | 1 |  |  |  | 1 |
| 3 | 3_95 | 36 | 1 | 1 |  |  |  | 1 |
| 3 | 3_96 | 35 | 2 | 0.75 | 0.25 |  |  | 1.6 |
| 3 | 3_97 | 84 | 1 | 1 |  |  |  | 1 |
| 4 | 4_1 | 37 | 2 | 0.95 | 0.05 |  |  | 1.1 |
| 4 | 4_10 | 49 | 1 | 1 |  |  |  | 1 |
| 4 | 4_100 | 63 | 2 | 0.95 | 0.05 |  |  | 1.1 |
| 4 | 4_105 | 53 | 1 | 1 |  |  |  | 1 |
| 4 | 4_106 | 12 | 1 | 1 |  |  |  | 1 |
| 4 | 4_107 | 41 | 1 | 1 |  |  |  | 1 |
| 4 | 4_11 | 50 | 1 | 1 |  |  |  | 1 |
| 4 | 4_110 | 64 | 2 | 0.95 | 0.05 |  |  | 1.1 |
| 4 | 4_111 | 13 | 1 | 1 |  |  |  | 1 |
| 4 | 4_123 | 60 | 1 | 1 |  |  |  | 1 |
| 4 | 4_125 | 97 | 1 | 1 |  |  |  | 1 |
| 4 | 4_126 | 37 | 1 | 1 |  |  |  | 1 |
| 4 | 4_127 | 12 | 1 | 1 |  |  |  | 1 |
| 4 | 4_128 | 72 | 1 | 1 |  |  |  | 1 |
| 4 | 4_15 | 81 | 1 | 1 |  |  |  | 1 |
| 4 | 4_18 | 37 | 1 | 1 |  |  |  | 1 |
| 4 | 4_25 | 71 | 1 | 1 |  |  |  | 1 |
| 4 | 4_27 | 48 | 1 | 1 |  |  |  | 1 |
| 4 | 4_28 | 56 | 1 | 1 |  |  |  | 1 |
| 4 | 4_3 | 67 | 1 | 1 |  |  |  | 1 |
| 4 | 4_33 | 66 | 2 | 0.95 | 0.05 |  |  | 1.1 |
| 4 | 4_34 | 32 | 1 | 1 |  |  |  | 1 |
| 4 | 4_36 | 25 | 1 | 1 |  |  |  | 1 |
| 4 | 4_37 | 69 | 1 | 1 |  |  |  | 1 |
| 4 | 4_38 | 61 | 1 | 1 |  |  |  | 1 |
| 4 | 4_39 | 68 | 1 | 1 |  |  |  | 1 |
| 4 | 4_40 | 82 | 1 | 1 |  |  |  | 1 |
| 4 | 4_44 | 37 | 1 | 1 |  |  |  | 1 |
| 4 | 4_45 | 22 | 1 | 1 |  |  |  | 1 |
| 4 | 4_47 | 92 | 1 | 1 |  |  |  | 1 |
| 4 | 4_53 | 31 | 1 | 1 |  |  |  | 1 |
| 4 | 4_54 | 44 | 1 | 1 |  |  |  | 1 |
| 4 | 4_55 | 43 | 1 | 1 |  |  |  | 1 |
| 4 | 4_56 | 55 | 1 | 1 |  |  |  | 1 |
| 4 | 4_59 | 53 | 1 | 1 |  |  |  | 1 |
| 4 | 4_60 | 36 | 1 | 1 |  |  |  | 1 |
| 4 | 4_61 | 10 | 1 | 1 |  |  |  | 1 |
| 4 | 4_63 | 59 | 1 | 1 |  |  |  | 1 |
| 4 | 4_65 | 137 | 1 | 1 |  |  |  | 1 |
| 4 | 4_66 | 25 | 1 | 1 |  |  |  | 1 |
| 4 | 4_67 | 80 | 1 | 1 |  |  |  | 1 |
| 4 | 4_68 | 31 | 1 | 1 |  |  |  | 1 |
| 4 | 4_69 | 63 | 1 | 1 |  |  |  | 1 |
| 4 | 4_7 | 59 | 1 | 1 |  |  |  | 1 |
| 4 | 4_70 | 40 | 1 | 1 |  |  |  | 1 |
| 4 | 4_72 | 77 | 1 | 1 |  |  |  | 1 |
| 4 | 4_75 | 15 | 1 | 1 |  |  |  | 1 |
| 4 | 4_78 | 29 | 1 | 1 |  |  |  | 1 |
| 4 | 4_79 | 72 | 1 | 1 |  |  |  | 1 |
| 4 | 4_8 | 46 | 1 | 1 |  |  |  | 1 |
| 4 | 4_80 | 48 | 1 | 1 |  |  |  | 1 |
| 4 | 4_84 | 56 | 1 | 1 |  |  |  | 1 |
| 4 | 4_87 | 58 | 2 | 0.6 | 0.4 |  |  | 1.92 |
| 4 | 4_89 | 25 | 1 | 1 |  |  |  | 1 |
| 4 | 4_90 | 61 | 1 | 1 |  |  |  | 1 |
| 4 | 4_91 | 36 | 1 | 1 |  |  |  | 1 |
| 4 | 4_92 | 87 | 1 | 1 |  |  |  | 1 |
| 4 | 4_94 | 62 | 1 | 1 |  |  |  | 1 |
| 4 | 4_96 | 77 | 1 | 1 |  |  |  | 1 |
| 4 | 4_97 | 80 | 1 | 1 |  |  |  | 1 |
| 4 | 4_98 | 38 | 1 | 1 |  |  |  | 1 |
| 4 | 4_99 | 65 | 2 | 0.95 | 0.05 |  |  | 1.1 |
